# Integrative analysis of large scale transcriptome data draws a comprehensive functional landscape of *Phaeodactylum tricornutum* genome and evolutionary origin of diatoms

**DOI:** 10.1101/176024

**Authors:** Achal Rastogi, Uma Maheswari, Richard G. Dorrell, Florian Maumus, Fabio Rocha Jimenez Vieira, Adam Kustka, James McCarthy, Andy E. Allen, Paul Kersey, Chris Bowler, Leila Tirichine

## Abstract

Diatoms are one of the most successful and ecologically important groups of eukaryotic phytoplankton in the modern ocean. Deciphering their genomes is a key step towards better understanding of their biological innovations, evolutionary origins, and ecological underpinnings. Here, we have used 90 RNA-Seq datasets from different growth conditions combined with published expressed sequence tags and protein sequences from multiple taxa to explore the genome of the model diatom *Phaeodactylum tricornutum,* and introduce 1,489 novel genes. The new annotation additionally permitted the discovery for the first time of extensive alternative splicing (AS) in diatoms, including intron retention and exon skipping which increases the diversity of transcripts to regulate gene expression in response to nutrient limitations. In addition, we have used up-to-date reference sequence libraries to dissect the taxonomic origins of diatom genomes. We show that the *P. tricornutum* genome is replete in lineage-specific genes, with up to 47% of the gene models present only possessing orthologues in other stramenopile groups. Finally, we have performed a comprehensive *de novo* annotation of repetitive elements showing novel classes of TEs such as SINE, MITE, LINE and TRIM/LARD. This work provides a solid foundation for future studies of diatom gene function, evolution and ecology.

## Introduction

Diatoms are one of the most important and abundant photosynthetic micro-eukaryotes, and contribute annually about 40% of marine primary productivity and 20% of global carbon fixation ^1^. Marine diatoms are highly diverse and span a wide range of latitudes, from tropical to polar regions. The diversity of planktonic diatoms was recently estimated using metabarcoding to be around 4,748 operational taxonomic units (OTUs) ^2,3^. In addition to performing key biogeochemical functions ^4^, marine diatoms are also important for human society, being the anchor of marine food webs, and providing high value compounds for pharmaceutical, cosmetic and industrial applications ^4^. A deeper understanding of their genomes can therefore provide key insights into their ecology, evolution, and biology.

Complete genome sequences for seven diatom species have been published ^5,6^, starting with the centric and pennate diatoms *Phaeodactylum tricornutum*^7^ and *Thalassiosira pseudonana,* respectively ^8^. Analysis of the *P. tricornutum* genome revealed an evolutionarily chimeric signal with genes apparently derived from red and green algal sources as well as the endosymbiotic host, and from a range of bacteria by lateral gene transfers ^7,9,10^, although the exact contributions of different donors to the *P. tricornutum* genome remains debated ^6,11,12^, alongside vertically inherited and group-specific genes. Such a diversity of genes has likely provided *P. tricornutum* and diatoms in general with a high degree of metabolic flexibility that has played a major role in determining their success in contemporary oceans.

The availability of a sequenced genome for *P. tricornutum* has also opened the gate for functional genomics studies, e.g., using Gateway, RNAi, CRISPR, TALEN and conjugation, which have revealed novel proteins and metabolic capabilities such as the urea cycle, proteins important for iron acquisition, cell cycle progression, lipid metabolism for biofuel production, as well as red and far/red light sensing^5^. Deeper understanding of the ecology and success of diatoms and a thorough dissection of the gene repertoire of *P. tricornutum* will however require better information regarding genome composition and gene structure. The first draft of the *P. tricornutum* genome (Phatr1), based on Sanger sequencing, was released in 2005, and contained 588 genome scaffolds totalling 31 Mb. The draft genome was further re-annotated and released as Phatr2 in 2008 with an improved assembly condensed into 33 scaffolds and 55 unmapped sequences which could not be assigned to any of the mapped chromosomes (denoted Phatr2 bottom drawer) ^7^, although many of the gene models contained within this annotation remained incomplete ^13^. Subsequently, sequencing technologies have evolved and RNA-Seq has been established as a gold standard for transcriptome investigation, and knowledge has advanced rapidly concerning the roles of DNA methylation and histone modifications on transcriptional regulation. Therefore, we exploited a large set of RNA-Seq reads derived from cells grown in different conditions, in combination with expressed sequence tags (ESTs) ^14^ and protein sequences (UniProt), as well as histone post-translational modifications and DNA methylation ^15,16^ to re-annotate the *P. tricornutum* genome. This allowed the identification of a significant number of novel transcripts including reverse transcriptase (Rv) with additional protein domains suggesting a domestication from the host of Rv domains that were used for genetic innovation by the host. Novel classes of transposable elements (TEs) were revealed in this annotation including MITE, SINE and LINE elements. We further identified extensive alternative splicing (AS) involved in regulation of gene expression in response to nutrient starvation suggesting that AS is likely to be used by diatoms to cope with environmental changes. We report a conserved epigenetic code, providing the host with different chromatin states involved in transcriptional regulation of genes and TEs. Finally, our work dissected the proposed complex chimeric nature of diatom genomes demonstrating the transfer of green, red and bacterial genes into diatoms, using the greatly expanded genomic and transcriptomic reference libraries that have become available across the tree of life since the publication of the initial genome ^10,17^. This resource was released as *Phaeodactylum tricornutum* annotation 3 (Phatr3) and is available to the community on the Ensembl portal (http://protists.ensembl.org/Phaeodactylum_tricornutum/Info/Index).

## Results and Discussion

### Structural re-annotation of *P. tricornutum* genome reveals numerous new gene models

To generate a new annotation of the *P. tricornutum* nuclear genome, our approach combined high-throughput RNA sequencing data (RNA-Seq) along with ESTs and protein sequences. Several mapping pipelines (see Methods) were used, allowing the prediction of 12,233 gene models with an average gene length of 1,624 bp and ∼1.7 exons per gene. The predicted Phatr3 gene models were then compared to the Phatr2 gene models (http://genome.jgi.doe.gov/Phatr2/Phatr2.home.html) and their structural differences can be grouped into the following categories (Table 1, Table S1):

**Table 1.**
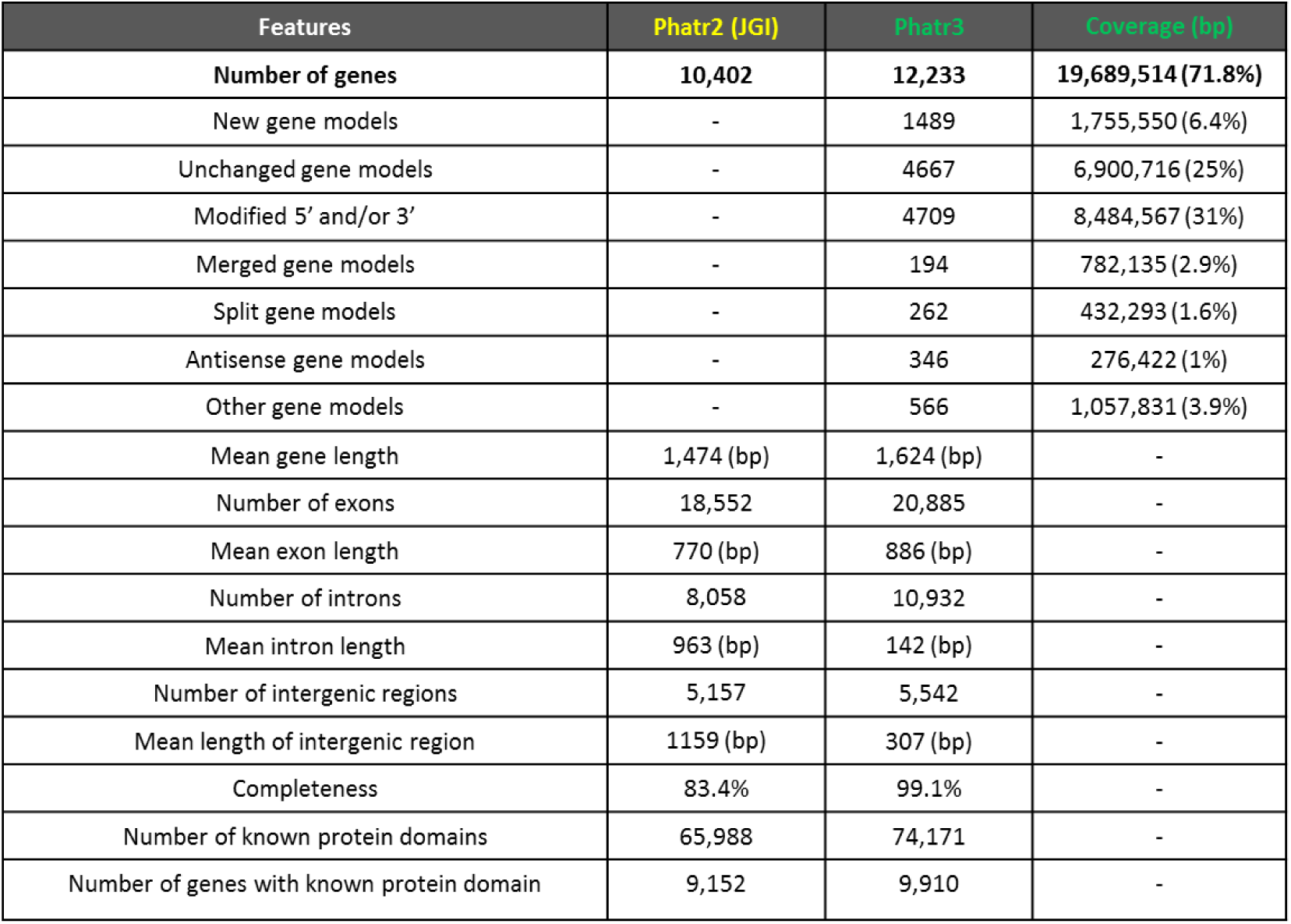
Comparison of Phatr3 and Phatr2 annotations. The table presents a summary of Phatr3 and Phatr2 gene comparison statistics. In case of Phatr2-JGI gene models, only filtered models have been considered in each case. The number of genes, in each category, and their corresponding coverage of the genome (in base-pairs) is given. Completeness here refers to the percentage of gene models found to contain both start and stop codons.

1. **New gene models,** genes that are newly discovered and are not present in Phatr2 gene annotations.
2. **Unchanged gene models,** gene whose structural annotation remains the same as in Phatr2.
3. **Modified gene models,** genes whose structural annotation has a different 5’ end, 3’end or both 5’ and 3’ ends with respect to Phatr2. Thirty of these genes with a different N terminus in Phatr3 compared to Phatr2 were validated by their presence within a previously constructed multi-gene reference dataset of aligned plastid-targeted proteins that are well conserved across ochrophyte lineages (File S1, panel A) ^10^. The N-terminus identified by the Phatr3 gene model in each alignment broadly matches the N-termini identified for orthologous sequences from other ochrophytes (File S1, panel B). Furthermore, RT-PCR analysis of six genes within this dataset amplified genes with product length predicted by Phatr3 (File S1, panel C).
4. **Merged gene models,** genes that are formed by merging two or more Phatr2 genes into one Phatr3 gene model. Such examples include proteins (e.g., Phatr3_EG02340, Phatr3_EG02341) that merge two domains which are often found together in several other organisms (e.g., a calcium binding EF-hand and a protein kinase). Likewise, a response regulator domain was found to merge with histidine kinase (Phatr3_EG02387), which both are part of the two-component regulatory system widely used by living organisms to sense and respond to changes in their environment ^18^. Several other examples of merged protein domains of the same pathway can be found in Table S1.
5. **Split gene models,** genes that are formed by splitting one Phatr2 gene into two Phatr3 gene models.
6. **Antisense gene models,** genes that are found localized on the antisense strand of previously annotated Phatr2 genes.
7. **Others,** 566 genes which do not fall into any of the categories above. These genes require manual curation which can be achieved through the Web Apollo Portal we implemented to improve the Phatr3 genome annotation. Since the length of 56 genes in the Phatr3 repertoire is less than 100 bp, we only considered 12,177 genes for further functional analysis (Table S1).

### Assessment of the conservation and complex evolutionary origin of *P. tricornutum* genome

In light of the recent availability of numerous genome and transcriptome sequences from many under-sampled taxa (e.g., red algae) through resources such as the Marine Microbial Eukaryote Transcriptome Sequencing Project ^10,17^, we wished to update and re-dissect the proposed complex chimeric nature of the *P. tricornutum* genome. We first aimed to assess the conservation of the *P. tricornutum* proteome across various taxonomic categories, which we grouped together based on recent published phylogenies and taxonomic reviews (File S2)^10,19^. From this analysis, a total of 9,008 (74.0%) of the genes within the *P. tricornutum* genome were found shared with at least one other group within the tree of life. This is substantially greater than the ∼ 60% of *P. tricornutum* genes previously identified to have orthologues in other groups ^7^, underlining the importance of dataset size and taxonomic sampling when considering gene conservation ^11^. Up to 251 different conservation patterns were identified across the entire genome, thirteen of which each accounted for 100 genes or more (Fig. 1). Many of the genes were found to have broad distributions across the tree of life, with 4,543 genes (37.3%) found in at least five of the nine groups considered. These included 203 genes (1.7%) in all groups studied, and 1074 (8.8%) in all groups except viruses (Fig. 1; Table S2). A further 1,188 genes (9.8%) were universally found across all eukaryotic groups but neither in prokaryotes nor viruses, hence might constitute eukaryote-specific genes (Fig. 1; Table S2).

**Figure 1.**
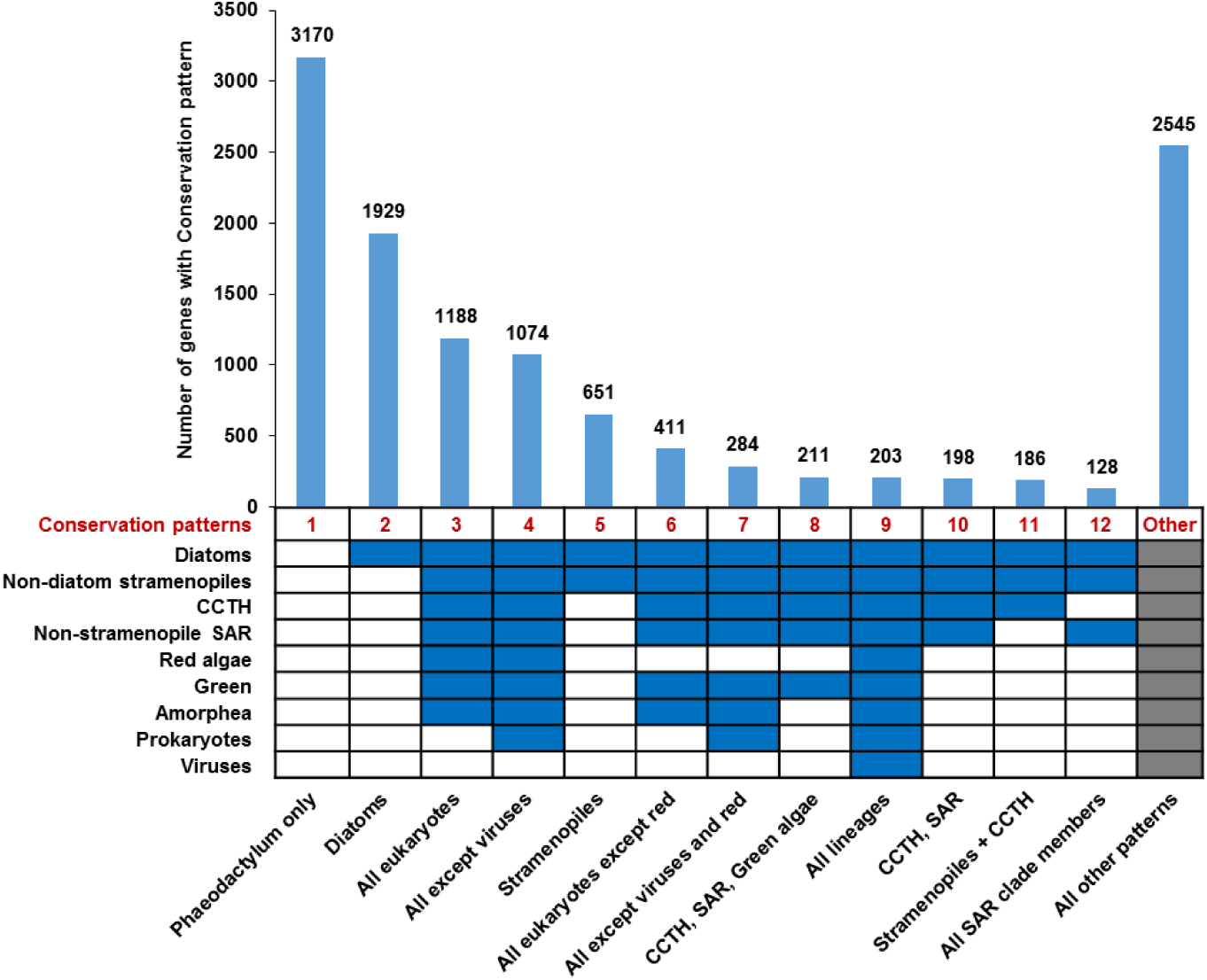
Conserved and group-specific genes in *P. tricornutum*. The chart shows the numbers of different Phatr3 genes shared with different combinations of groups. "SAR" refers to the combined clade of stramenopiles, alveolates and rhizaria; "CCTH" the combined clade of cryptomonads, centrohelids, haptophytes and telonemids; and "amorphea" the combined clade of opisthokonts, amoebozoa and excavates^19^. Below the bar-plot, a heatmap gives an overview of twelve conservation patterns (shown as columns), each of which account for at least 100 genes in the Phatr3 genome. Blue cells indicate that orthologues are detected within two or more sub-categories within the corresponding lineage (or are detected at all within viruses, for which only one sub-category was considered), white cells that orthologues are detected in fewer than two sub-categories (or not detected at all for viruses), and grey cells that either conservation pattern was permitted. Above, the bar-plot shows the number of genes associated with each conservation pattern. A similar plot for new gene models only is shown in Fig. S1.

We still found, with the expanded dataset, that many genes within the *P. tricornutum* genome have limited evolutionary conservation, with 5750 genes (47.2%) having originated within the recent vertical history of the stramenopile lineage. A total of 3,170 genes (26.0%) were found to be specific to *P. tricornutum*, 1,929 were only shared between *P. tricornutum* and other diatoms (15.8%), and 651 were only shared with diatoms and other stramenopiles (5.3%) (Fig. 1). We found only limited evidence for genes that were not shared between *Phaeodactylum* and other diatoms, but were shared with other groups (410 genes, 3.4%), or for genes that were not shared between *Phaeodactylum* and other stramenopiles but were shared with other groups (242 genes, 2.0%), suggesting largely vertical recent inheritance of the *Phaeodactylum* genome (Table S2).

Further, we wished to determine whether the 1,489 novel genes uncovered by Phatr3 differ in terms of evolutionary conservation to those previously identified. While many of the novel genes are specific to *P. tricornutum* (864 genes; 58.0%; Fig. S1) or are limited to diatoms (222 genes; 14.9%; Fig. S1), 44 genes (13.6%) are shared with at least five other groups, and 4 novel genes are shared with all nine groups considered (Fig. S1), including a UvrD-like DNA helicase containing protein (Phatr3_EG00261), CTP biosynthetic process (Phatr3_EG00931), telomere recombination (Phatr3_J11434) genes and a high motility group protein (Phatr3_J1241), confirming that many of these genes are likely to have important biological functions.

In our second analysis, we aimed to reassess the evolutionary origins of the *P. tricornutum* genome. In particular, we wished to validate the presence of genes derived from green algae and from prokaryotes, which have previously been controversial ^7,9,11^, and identify whether different predicted gene transfer events occurred specifically in *P. tricornutum*, or are more ancient events, occurring prior to the radiation of pennate diatoms, all extant diatoms, stramenopiles, or previously.

#### Prokaryotic genes

Across the entire dataset, 584 genes yielded top BLAST hits against prokaryotes (Table S3), which is similar to the number of prokaryotic genes (587) identified in the initial publication of the *P. tricornutum* genome ^7^. Similarly to the initial genome publication, the prokaryotic sub-category that produced the most top hits (235) was the proteobacteria (Table S3; Fig. S2A). Nine other sub-categories (cyanobacteria, firmicutes, chlorobi, archaea, actinobacteria, chlamydiae, chloroflexi, the *Deinococcus-Thermus* clade, and planctomycetes) contributed more than ten hits each (Fig. S2A; Table S3). The 15 gene transfers involving members of the *Deinococcus-Thermus* clade are of particular interest, as this lineage has not previously been reported to have specifically exchanged genes with an ancestor of *P. tricornutum*^7^.

We considered whether the prokaryotic genes present in the *P. tricornutum* genome are recent acquisitions (e.g., species-specific), or occurred at earlier points in the evolution of diatom lineages. This was not possible in the initial genome, for which the only other available diatom genome was for the centric species *T. pseudonana*^7,8^. For this, we performed an analysis in which we serially removed the closest relatives of *P. tricornutum* in our sequence library (which includes seven complete diatom genomes and transcriptomes for a further 92 diatom species available through MMETSP)^11,36^, and assessed the number of prokaryotic genes that could be identified in each analysis. Twenty two of the prokaryotic genes were identifiable with the full dataset (hence were specifically acquired by *P. tricornutum* following its divergence from other diatoms), 69 were identifiable with a full dataset excluding pennate diatoms (hence were presumably acquired within the evolutionary history of the pennate lineage) and 202 were identifiable with the full dataset excluding all diatoms (hence were acquired during the early evolution of diatom lineages, prior to the division of pennate lineages from their closest relatives within the polar centric diatoms), with the remaining 291 showing more ancient origins (Table S3). Thus multiple gene transfer events involving prokaryotes have occurred progressively through the evolution of ancestors of *P. tricornutum.*

#### Red algal genes

Across the entire dataset, 459 genes produced BLAST top hits against members of the red algae, consistent with the red algal ancestry of the diatom plastid ^4,20^ (Fig. 2A). This is broadly equivalent to the number of red genes identified in previous studies of diatom plastids ^21^. The two sub-categories with the greatest contributions to these genes were the Porphyridiophytes (150 genes) and Bangiophytes/Florideophytes (147 genes) (Fig. S2B; Table S3). A total of 353 of the red algal genes were identified following removal of all ochrophyte sequences from the dataset, with only a further 28 identified following the removal of aplastidic stramenopile groups (oomycetes, labyrinthulomycetes, and slopalinids) and a further 25 identified following the removal of the two remaining SAR clade groups (ciliates, and aplastidic rhizaria) considered (Fig. 2B; Table S3). The limited number of genes of red algal origin identified within aplastidic SAR clade members supports a late acquisition of a red algal plastid by a common ancestor of all ochrophytes, following their divergence from oomycetes ^10,22^.

**Figure 2.**
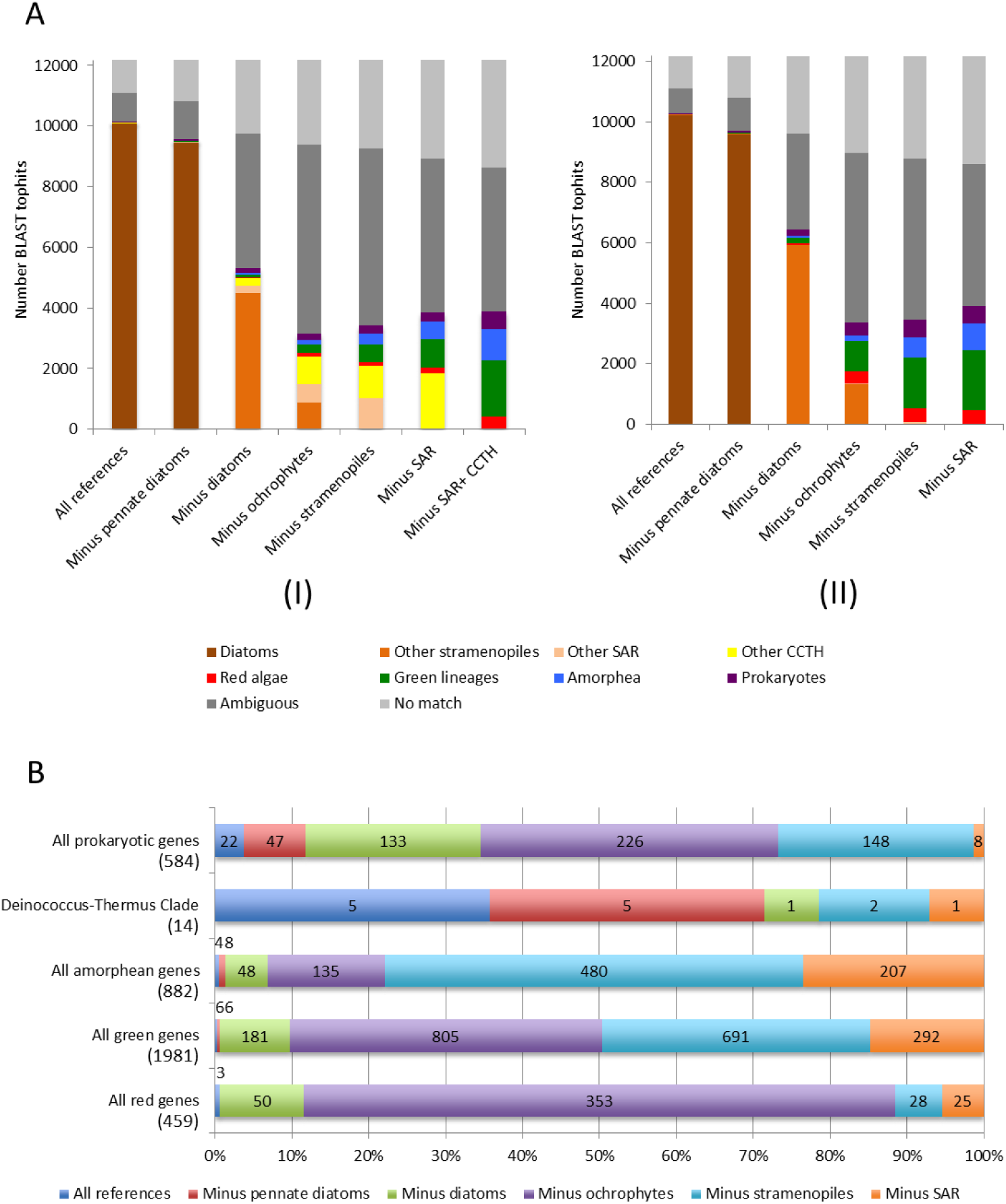
BLAST top hit analysis of *P. tricornutum* genes. (A) Comparison of the number of genes of unambiguous taxonomic origin, identified by BLAST top hit analysis of the complete Phatr3 protein annotations against six different reference libraries, constructed using UniRef, jgi genomes and other transcriptome libraries, and six different patterns of taxon inclusion: a library containing all sequences from the tree of life; all sequences except those from pennate diatoms; all except those from diatoms; all except those from stramenopiles; all except those from SAR clade members; and all except those from SAR and CCTH clade members. The right-hand graph (II) shows similar values calculated for reference libraries from which all algae with a suspected history of secondary endosymbiosis other than ochrophytes were removed. Note that in this case a separate value for a reference library excluding SAR and CCTH clade taxa is not provided, as all it is not known when secondary endosymbioses arose within CCTH clade lineages ^19^, hence all CCTH clade taxa were excluded from the analysis. (B) Comparison of the proportion of genes that yield top hits against five different categories (all prokaryotes, the *Deinococcus-Thermus* clade only, all red algae, and green groups, and all amorphea) following the removal of different groups from the complex algae-free dataset. Total number of genes within each category is indicated beneath it in round brackets. Data labels for each value are provided in the corresponding segment; where the segment is too small to accommodate the corresponding value, the value is placed outside the chart, and shaded to match the color of the segment. The largest value recorded for each gene category corresponds to the most probable evolutionary time point at which genes were acquired; for example, the largest number of genes of red algal affinity were recovered following the removal of all algae with plastids of secondary or higher endosymbiotic derivation from the reference dataset, indicating a large-scale donation of red algal genes into algae with plastids of secondary or higher endosymbiotic derivation.

#### Green genes

A total of 1,981 genes generated top BLAST hits from members of the green group (green algae and plants). This is similar in size to the number of green genes (>1700) identified in previous studies of the origins of diatom groups ^21^, and could be consistent with large scale gene transfer between diatom ancestors and green algae (Table S3). Some of these genes may be misidentified genes of red algal origin, as has been discussed elsewhere ^11,12^; however, we believe that many are genuinely of green origin, for two reasons. Firstly, compared to previous phylogenomic studies of diatom genomes, our reference library contains a much larger amount of red algal sequence information, including five complete genomes, and large-scale transcriptomes for a further twelve red algal species (Table S4) ^10^. Up to 685 of the identified green genes had orthologues (as confirmed by the reciprocal best-hit (RbH) analysis) in two or more red sub-categories, 314 had identified orthologues in two or more subcategories each of red algae, green groups, amorphea (opisthokonts, amoebozoa and excavates)^19^ and prokaryotes, and 222 had identified orthologues in all five of the red sub-categories and all eleven of the green sub-categories considered (Fig. S3A). We saw no difference in the representation of red and green algal sub-categories in genes with annotated red or green origin (Fig. S3B).

Secondly, green gene transfers appear to have occurred at a different time point to the red algal gene transfers. Although the largest number of putative green genes (805) were identified with the dataset from which all ochrophyte groups were removed (Fig. 2B), nearly as many (691) were identified following the removal of aplastidic stramenopiles from the dataset (Fig. 2A). This contrasts to the situation for red genes (which were overwhelmingly identified following the removal of all ochrophyte sequences from the library, as discussed above; Fig. 2A), and might potentially indicate two distinct gene transfer events between the green algal and stramenopile lineages: an early transfer of green genes in an ancestral stramenopile ancestor, and a subsequent transfer of green genes into an ochrophyte plastid, possibly concomitant with or mediated via the acquisition of a chimeric ochrophyte plastid ^10^. In summary, our data therefore supports previous findings ^10,21^ of gene transfers between an ancestor of stramenopiles and one or more groups of chlorophyte algae. More broadly, the presence of green, red and prokaryotic genes in the *P. tricornutum* genome, which appear to have arisen at different points in its evolutionary history, confirms that it is an evolutionary mosaic, reflects examples of inferred gene transfer events in other major eukaryotic algal lineages ^10,23,24^, and underlines the significance of progressive horizontal gene transfer in the evolution and diversification of modern algae ^25^.

### Update of the functional annotation of the *P. tricornutum* proteome

We next performed gene ontology analysis of the Phatr3 genes using UniProt-GOA (UniProt release 2015_03) and implemented a detailed analysis of their functional domain architecture using DAMA and CLADE^26,27^. From the analysis we found 12,092 genes (∼99%) with known functional domains, which can be grouped into 5,021 gene families. Among all, the largest gene families are with genes containing reverse transcriptase (Rv) domains (169 genes), RNase H domain (154 genes) and Integrase domain (132 genes). These functional domains often co-localize and are associated with transposable elements. Apart from the latter, protein kinase gene family (115 genes) is also abundant (Table S1) within *P. tricornutum*.

Interesting domain architectures were found among the genes that contain either Rv, RNase or H domain alone or together with additional protein domains, including 41 genes possessing Chomo (CHromatin Organization Modifier) domain or 45 cyclins, N/C-terminal domains (Table S1). The large number of sequences encoding reverse transcriptase domains in the *P. tricornutum* genome reflects previous studies that suggest these proteins are highly abundant and transcriptionally active in diatoms, implicating a possible role in their evolution and adaptation to contemporary environments ^28^. The presence of additional protein domains supports previous studies ^29^ which suggest that Rv domain-containing proteins might have originated from domesticated retrotransposons that evolved different functions via acquisition of various N- and C-terminal extensions.

We further aimed to determine whether genes with different levels of conservation, as determined by our analysis (Table S2), have different functional properties in the *P. tricornutum* genome. We performed GO enrichment analysis on four different biological categories of genes, as defined by the presence or absence of orthologues in other lineages by RbH analysis (Fig. 1). These were: the 3170 genes that are specific to *P. tricornutum* (Pt-specific genes), the 1929 genes that are uniquely shared with other diatoms (diatom-specific genes), the 1188 genes that are shared across all eukaryotic groups, and the 203 genes that are shared with all other eukaryotic groups and with prokaryotes (Fig. 1; Fig S4; Table S5). Interestingly, a high number of Pt-specific genes encode the DNA integration GO category, which may indicate a permissive way for integration of genetic material from diverse sources, thus creating novel genetic diversity. Of note, one Pt-specific gene (Phatr3_J49482) encodes a Tir chaperone protein which functions in type III secretion system involved in pathogenicity, although there is no evidence so far for the pathogenic potential of *P. tricornutum.* Regulation of transcription is one of the important functional categories of diatom-specific genes, perhaps reflecting differences in promoter architecture compared to other eukaryotes such as animals and plants. A large eukaryotic gene family shared with prokaryotes encodes oxidation-reduction processes that may have relevance for maintaining the homeostasis of unicellular organisms for an efficient metabolism.

Next, we used ASAFind and HECTAR to predict the sub-cellular targeting of the Phatr3 proteome (Table S6). Across Phatr3, 3196 proteins (26.3% total; Fig. S5A) were predicted to have a targeting sequence of some description by ASAFind, and 4067 proteins (33.3%; Fig. S5B) were predicted to have a targeting sequence by HECTAR. We then compared the Phatr3 proteome targeting predictions with the new Phatr3 gene models and Phatr2 JGI gene models. A greater proportion of the Phatr3 genes, including the new gene models, contain complete N-termini than Phatr2 (Fig. S5), reflecting that Phatr3 genes are better annotated structurally (File S1). Using both analyses, non-trivial numbers of proteins were predicted to have targeting predictions to individual cellular organelles, namely the plastid, endomembrane system or mitochondria (Fig. S5; Table S6). Several of the new gene models with defined targeting preferences had functions consistent with their localization: for example, we identified through ASAFind a plastid-targeted serine acetyltransferase (Phatr3_EGO1815), which forms an essential component of plastid cysteine synthesis pathways in ochrophytes ^10,30^, and a plastid-targeted betahydroxyacyl-ACP dehydratase protein (Phatr3_draftJ1143), which is consistent with the plastidial localization of fatty acid synthesis pathways in diatoms ^10,31^. The presence of domains performing known biological processes with consistent subcellular localization predictions, confirms that many of the novel genes identified within the *P. tricornutum* genome possess specific biological functions.

We also considered the expression dynamics of each gene using quartile approach (where elements of 1^st^ quartile are considered to have genes with no or low expression, 2^nd^ quartile with low to moderate expression, 3^rd^ quartile with moderate to high expression, and 4^th^ quartile with very high expression). Most of the novel genes (∼70%) are expressed at below the median level inferred for all other genes in the genome (Fig. 3A) and are mostly specific to *P. tricornutum* (Fig. S1). We then compared chromatin marks associated with new versus unchanged Phatr3 gene models and found that the proportion of DNA methylated genes within new gene models (30%, 448 genes) was found to clearly delineate the proportion within unchanged gene models (9%, 4,667 genes) (Table S1). The majority of these are at least methylated in CG context, which is in line with previous work ^15^. Thus, along with boosting the functional content of the genome, newly discovered genes will certainly expand our capacity to understand the role of DNA methylation in the regulation of *P. tricornutum* molecular machinery. Broadly, among the 1,489 new genes, 1,360 (∼91%) genes are marked by at least one of the epigenetic modifications studied previously (DNA methylation, H3K27me3, H3K9me2, H3K9me3, H3K4me2 and H3K9_14Ac). Most of these genes (563 genes, 38%; Table S1, Fig 3B) are marked by chromatin modifications associated with active chromatin states (H3K4me2 and H3K9_14Ac). On the other hand, 407 genes (27%) are marked exclusively by repressive chromatin modifications (DNA methylation, H3K27me3, H3K9me2, H3K9me3), and 390 genes (26%) are co-marked by both active and repressive marks with multiple combinations (Table S1, Fig 3B). The co-localization effect of different chromatin-level modifications on these new genes regulates repressive, active and moderate states of expression of the genome (Fig 3B).

**Figure 3.**
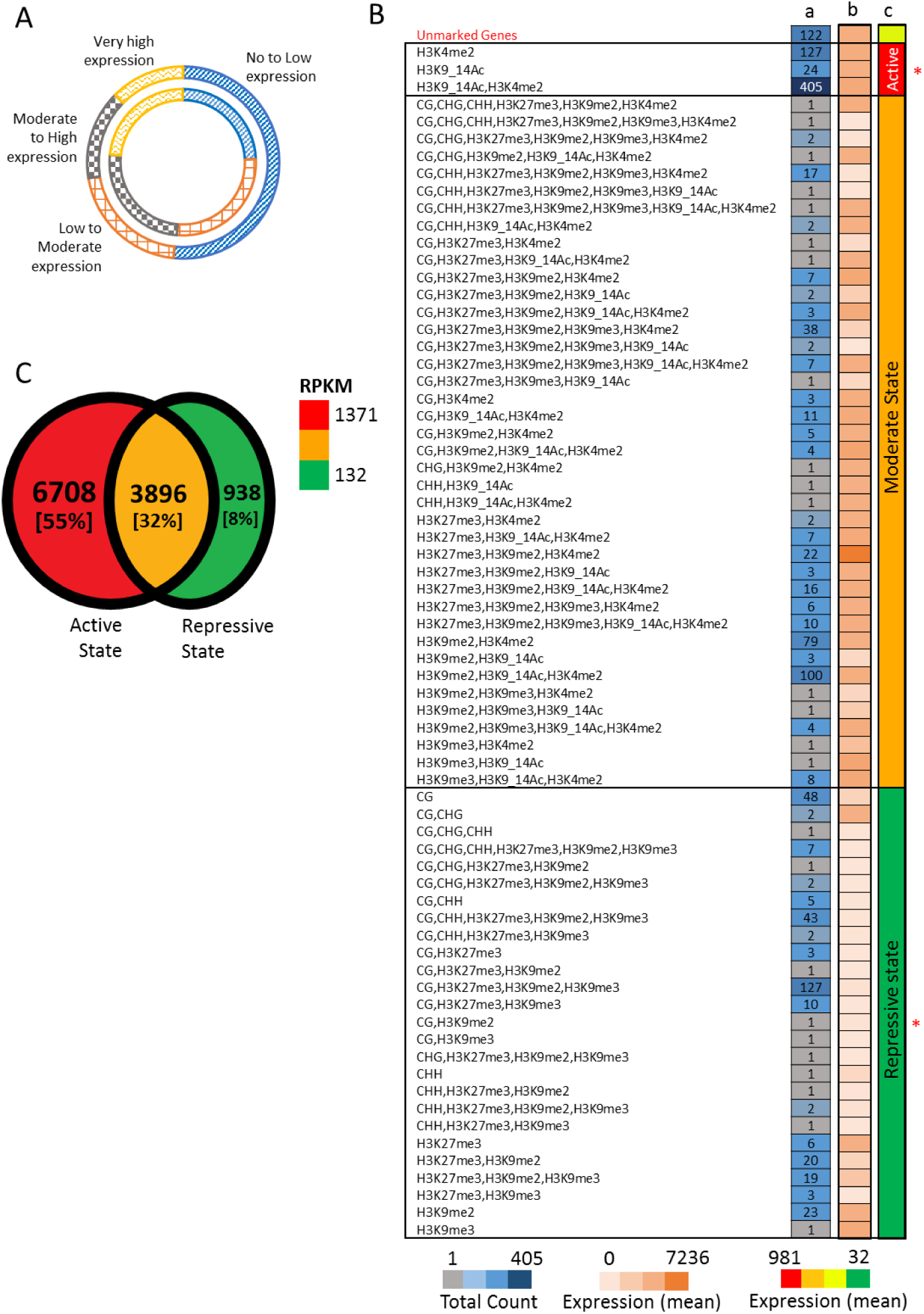
Dissecting the functional characteristics of Phatr3 proteome. (A) Comparison of the expression profile of the proportion of new genes (outer circle) with that of the proportion of unchanged gene models (inner circle). The expression profiling of all the Phatr3 genes are done based on quartile method, where first and last quartile reflects genes with no/low expression and very high expression, respectively. (B) and (C) summarize various epigenetic modification and regulatory effects on Phatr3 gene models and only new gene models in Phatr3. (B) Specifies the regulatory effects of the co-localization of different modifications/marks on the novel genes. The number of genes corresponding to each combination of co-localizing marks is provided in column a. The average expression of these genes is represented as a heat map in column b. Compared to the unmarked genes the regulatory effect of the co-localization of different epigenetic modifications is seen to maintain three chromatin states, represented with column c with few exceptions with expression that can be higher or lower than expected. This cases are likely due to the presence of additional active or repressive marks that were not studied. * indicates where the average expression (normalized DESeq counts) is significantly different (two sample T-test with unequal variance, P-value < 0.002), compared to the expression of unmarked genes. Out of 1,489 genes, 1,388 are considered in the analysis, as expression for the other 101 genes was not calculable without errors. (C) Different chromatin states maintained genome wide, based on the association of protein coding genes with repressive, active or both repressive and active chromatin modifiers (Repressive modifications: CG, CHG, CHH, H3K27me3, H3K9me2, H3K9me3; Active modifications: H3K4me2, H3K9_14Ac). Numbers and percentages (in square brackets) in the Venn diagram reflects the absolute number of genes and the relative percentage of the total Phatr3 genes.

Finally, we considered an update on the distribution of DNA methylation and post-translational modifications of histone H3 (PTMs) across the *P. tricornutum* genome, following previous work ^15,16^. Up to 11534 genes (∼95%) within Phatr3 were found to be either associated to the studied H3 PTMs or to DNA methylation. Most of the genes are preferentially labelled only by marks (6708 genes, ∼55%) associated with an active transcriptional state, such as acetylation (H3K9_14Ac) and/or H3K4me2 (Fig 3C; Table S1), whereas ∼8% genes are marked only by repressive modifications (DNA methylation, H3K27me3, H3K9me2 and H3K9me3) (Fig 3C; Table S1). A total of 3896 genes (∼32%) are marked by both active and repressive marks (Fig 3C; Table S1) suggesting a crosstalk between acetylation, H3K4me2 which are active marks and the remaining repressive marks inducing a combinatorial effect on gene expression. Overall, this analysis supports the conservation of a chromatin-level code, as previously reported in *P. tricornutum* ^15^, which is critical for transcriptional regulation of genomes and will be useful to decipher the role of epigenetics in underpinning the ecological success of diatoms.

### Intron retention is prominent in *P. tricornutum* and regulates genes under fluctuating environmental conditions

The functional and regulatory capacity of eukaryotic genomes is greatly influenced by alternative splicing of the precursor RNAs. Intron-retention (IR) is a major constituent of the alternative splicing code in plants and unicellular eukaryotes^32^, whereas exon-skipping (ES) is prominent within members of the metazoan clade; however, the broader evolutionary histories of both processes across the eukaryotes remains unclear ^33,34^. Therefore, and considering the position of diatoms in the tree of life, we investigated the nature and dynamics of both processes within the Phatr3 genome.

First, we mapped the distribution and dynamics of introns in the *P. tricornutum* genome. In total, we found 8646 introns (on average 0.7 introns per gene) with an average size of 142 bp, from which >99.8% include canonical splice-sites (Acceptor Sites: AG/CT and Donor sites: GT/AC). Up to 4014 (∼33%) of the Phatr3 genes were predicted to contain only one intron, while 1730 (∼14%) genes contain more than one intron and 6434 (∼53%) are predicted to be intron-less. The low density of introns and small intron size observed in *P. tricornutum* is similar to many unicellular eukaryotes, including the related diatoms *T. pseudonana* and *Fragilariopsis cylindrus* ^33,35,36^, which might mirror genome size, or enhance transcriptional efficiency or splicing accuracy in metabolically fluctuating environments ^37,38^. Notably, in *P. tricornutum* we found no difference between the average lengths of the first intron with respect to the others. This is similar to some species of the genus *Phytophthora* (Fig S6) and to what has been reported in *Schizosaccharomyces pombe* and *Aspergillus nidulans* ^39^. However, in most unicellular eukaryotes first introns are found to be significantly longer than non-first introns (Fig S6) ^39^. The functional consequences of this intron organization remain to be determined.

Next, we profiled alternative splicing events in *P. tricornutum* using RNA-Seq data generated in different growth and stress conditions (see Methods). From the 12177 Phatr3 gene models, 2924 (∼24%) genes are found to have introns that can be retained in more than 20% of the total experimental samples studied, while 2444 (∼20%) genes are observed to skip one or more exons in various samples. A total of 1335 (∼11%) genes are found to undergo both ES and IR, hence can perform alternative splicing (Fig 4A; Table S1). Like most unicellular eukaryotes and unlike metazoans, *P. tricornutum* shows a higher rate of IR than ES, supporting the hypothesis that ES has become more prevalent over the course of metazoan evolution ^32^. We then considered the expression dynamics of *P. tricornutum* genes that undergo IR or ES (Condition used: WT, Bio sample accession: SAMN06350643). Surprisingly, we found that genes that can undergo intron-retention are more highly expressed than genes that do not show alternative splicing (two sample t-test, P-value < 0.008, Fig 4B). This is in contrast to the situation in mammals in which intron-retention down-regulates the genes that are physiologically less relevant ^40^.

**Figure 4.**
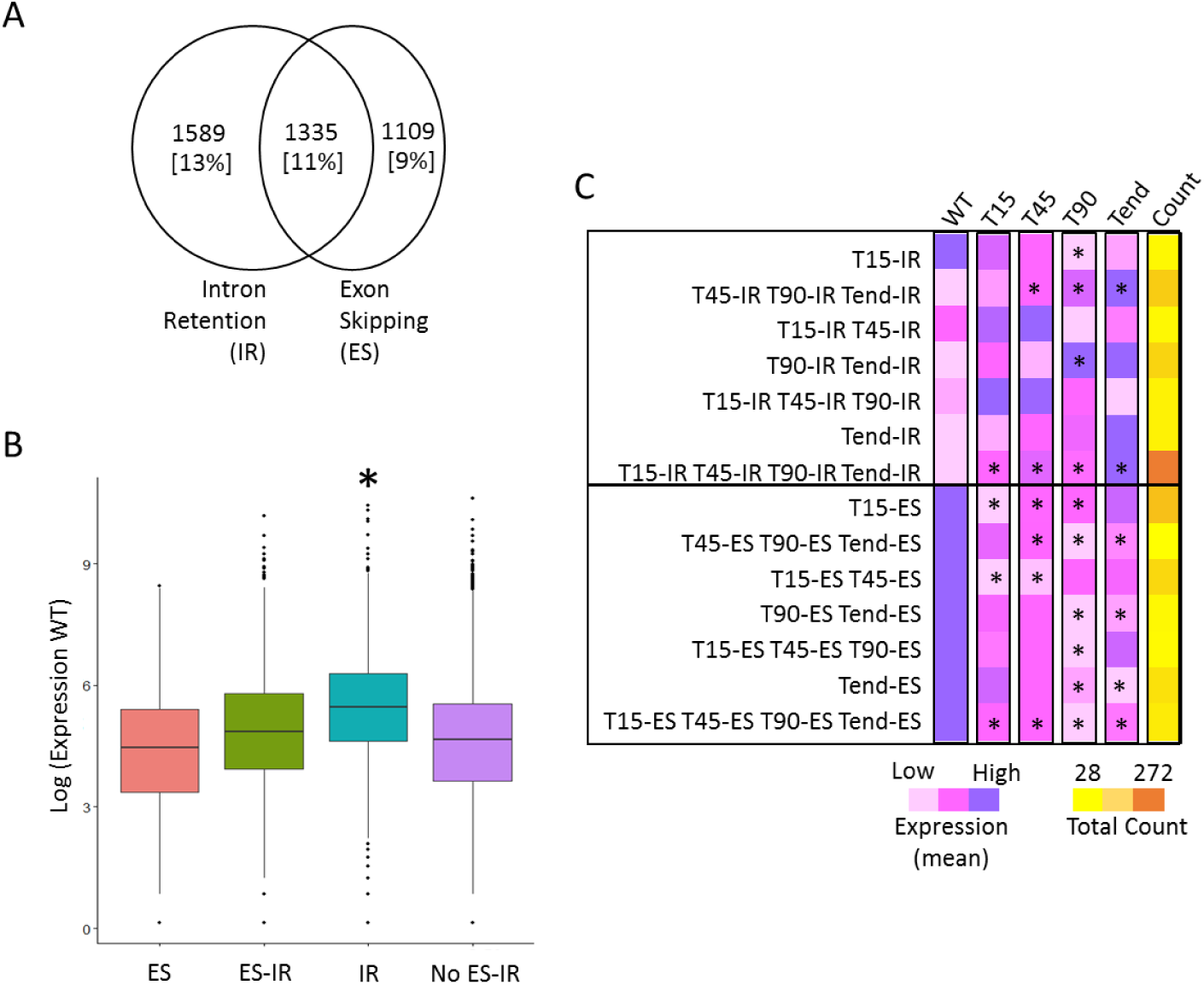
Alternative splicing in *P. tricornutum*. The Venn diagram depicts (A) the number of genes predicted to undergo alternative splicing in the context of intron retention (IR) and exon-skipping (ES). Numbers and percentages (in square brackets) in the Venn diagram reflect the absolute number of genes and the relative percentage out of the total Phatr3 genes. (B) The box-plot represents the differences in expression of genes corresponding to different functional categories, represented on x-axis. Y-axis represents the expression of the genes in the wild-type (WT) (Biosample accession: SAMN06350643), scaled to log scale. (C) The heat-map compares the average expression of genes exhibiting intron-retention/exon-skipping (IR/ES) at different time-points (T15, T45, T90 and Tend) in Nfree culture conditions. For example, T15-IR row compares the average expression of all the genes exhibiting intron retention (IR) are time T15 across all the time-points. The last column represents the heat-map based on the number of genes in each category indicated on left Y-axis. * indicates the level of significance as being significant (P-value < 0.05, two sample t-test with unequal variance) when the average expression of a particular sub-set is compared to the WT. Nfree in the figure denotes the culture condition with no source of nitrogen, WT denotes wild type or normal condition, T15 denotes RNAseq performed on cells sampled after 15 minutes of the culture, Similarly, T45, T90 and Tend denotes RNAseq libraries were prepared upon sampling the cells after 45 minutes, 90 minutes and 18 hours of culture, respectively.

To further assess the biological role of alternative splicing, we identified 1341 genes showing IR and 1099 genes with ES during an 18-hour time course under nitrogen-free growth conditions (see Methods). By comparing the expression levels and intron dynamics of each gene during the time course, we found a significant increase in the expression levels of genes undergoing intron retention (two sample t-test with unequal variance, P-value < 0.05) (Fig 4C; Table S1). For example, genes in which intron retention was observed from 45 minutes in the time course showed greater expression levels from this point onwards than in the immediate time period following the induction of the nitrogen starvation condition. In contrast, we observed a significant decrease in the expression of genes undergoing ES across different time-points of nitrogen-free growth (two sample t-test with unequal variance, P-value < 0.05) (Fig 4C). This indicates that the role of restructuring of genes via AS is non-trivial in maintaining the physiology of the cells under fluctuating environmental conditions.

We further examined the functions of the genes that are alternatively spliced under nitrogen starvation. We identified 81 GO categories which were significantly over-represented in either the IR or ES across all or different time-points (Table S7; Fig S7). Following the expression dynamics identified above, the GO categories over-represented in IR datasets show increased expression levels, and the GO categories in ES datasets diminished expression levels, from the point of induction over the length of the time-course. Many of the GO categories that show differential IR or ES dynamics have plausible functions in tolerating nitrogen starvation. For example, many of the genes that show enhanced intron-retention following 45 minutes of starvation are implicated in cell cycle regulation (e.g., functions in chromosome separation, histone modifications, or the mitotic cell-cycle checkpoint), which might correspond to an arrest of the cell cycle under nitrate limitation^41^. Similarly, genes implicated in catabolism and storage of cellular nitrogen pools (allantoin biosynthesis, glutamine biosynthesis, and glutamate catabolism) and nitrate and nitrite transport show enhanced intron-retention within 45 to 90 minutes of starvation induction^42^. Our data broadly suggest that ES, besides creating mRNA diversity, seems to be used for transcriptional regulation of specific genes under specific conditions in *P. tricornutum* and is likely to be widespread.

### Copia-type LTR makes up most of the TEs in the *P. tricornutum* genome

In the context of the Phatr3 re-annotation of *P. tricornutum* genome, we also revisited the annotation of repetitive elements in the genome assembly. In the current analysis, we applied a robust and *de novo* approach for the whole genome annotation of repeat sequences. Collectively, repeats were found to contribute ∼3.4 Mb (12%) of the assembly, including transposable elements (TEs), unclassified and tandem repeats, as well as fragments of host genes (Table 2). TEs are the dominant repetitive elements in *P. tricornutum* and represent 75% of the repeat set, i.e., 2.3 Mb as compared to 1.7 Mb in the previous TE annotation. By comparing the Phatr3 repertoire of TEs, including both large and small elements, with the previous TE annotations, 1988 (∼54%) TEs were found to be novel (Table S8).

**Table 2.**
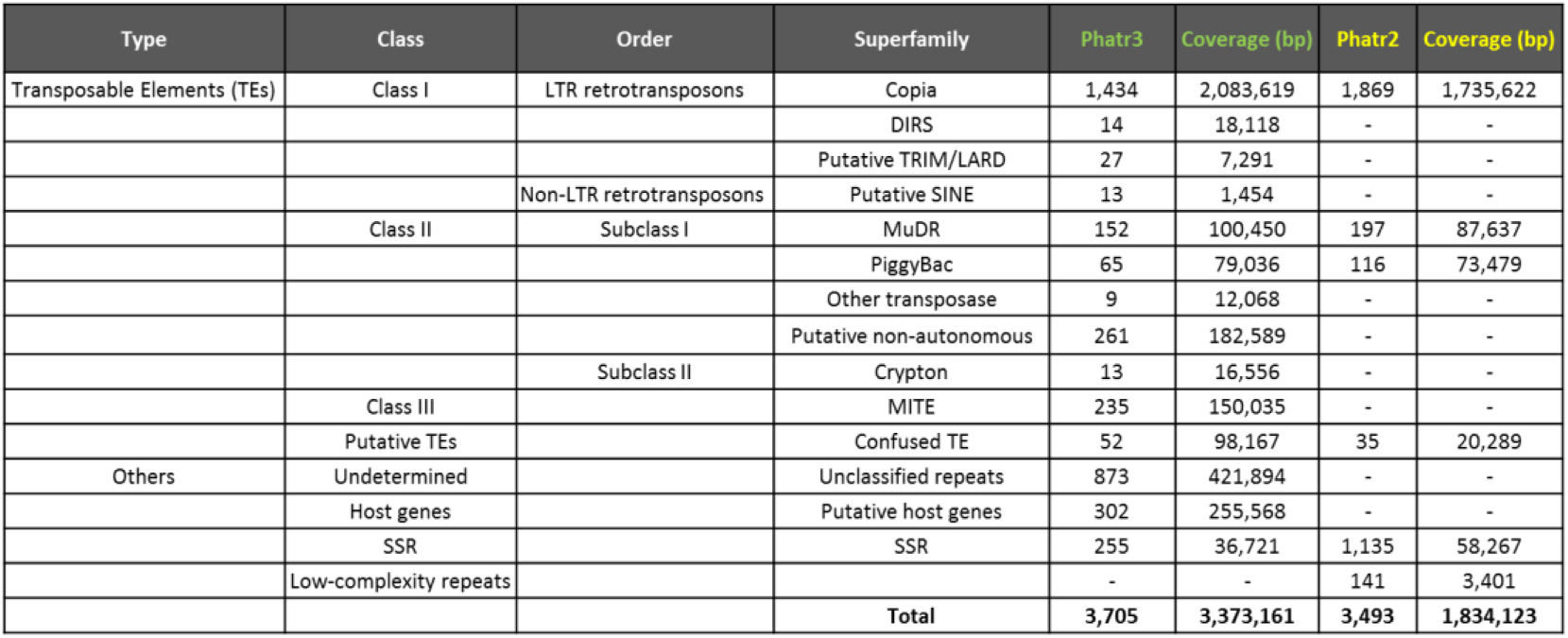
Composition of repetitive sequence content.

In line with previous analyses, Copia-type LTR retrotransposons (LTR-RTs) are the most abundant type of TEs, contributing over 55% of the repeat annotation, while Gypsy-type LTR-RTs remain undetected. This new TE annotation also reveals for the first time the presence of Crypton-type transposons in *P. tricornutum,* which are also found in fungi and multiple invertebrate groups ^43^. Previous examination of repetitive elements relied mainly on a library of manually curated TEs ^16,44^. In the present work, the de-novo annotation using current state-of-the-art approaches (see Methods) with all types of repeated elements yields more elements of the main classes of TEs in Phatr3. As a result, we detected more Copia-type elements, which cover approximately 200 kb of the genome. Furthermore, we annotated ∼183 kb of the genome that corresponds to copies of potential non-autonomous DNA transposons. Additionally, we detected for the first time Miniature inverted–repeat transposable elements (MITE) in a diatom which are known to be prevalent in plants and animals playing a major role in in genomes organization and species evolution.

Next, we considered the epigenetic marks and expression profiles associated with TEs within Phatr3. Consistent with previous reports ^15,16^, the majority (2790, ∼75%) of the Phatr3 TE repertoire is associated with one or other studied chromatin marks known to maintain either active (H3K4me2, H3K9_14Ac) and/or repressive states (DNA methylation, H3K27me3, H3K9me2, H3K9me3) of the genome (Fig S8A; Table S8). In contrast to coding regions, (including the new gene models), ∼50% (1845) of the TEs are marked solely by repressive marks (Fig S8A; Table S8), and only ∼7% (268) TEs are marked solely by active chromatin marks (Fig S8A; Table S8). A total of 677 TEs (∼18%) were found to be marked by both active and repressive marks (Fig S8A; Table S8). Profiling the chromatin landscape and DNA methylation specifically associated with new TEs revealed that 1,236 (∼62%) new TEs are found to be marked by at least one of the epigenetic marks, from which only 75 (∼6%), 25 (∼2%), 111 (∼9%), 58 (∼5%) and 62 (∼5%) are marked specifically by H3K27me3, H3K9me3, H3K9me2, H3K9-14Ac and H3K4me2, respectively (Fig S8B; Table S8). A total of 458 (∼23%) TEs were methylated, most of which (368; ∼80%) in a CG context only (Fig S8C; Table S8), while 19 (∼4%) TEs are specifically methylated in a CHH context, and 34 (∼7%) are found methylated by at least two sub-contexts of DNA methylation (CG, CHH and CHG) (Fig S8C; Table S8). As for genes, TEs marked by active PTMs of histones show high levels of expression while those that carry repressive marks display lower expression levels, and TEs with combinations of both marks are expressed at intermediate levels (Fig S8A, S8D). TEs that are methylated specifically in CG context are typically expressed at lower levels compared to those specifically methylated in CHG or CHH contexts (Fig S8E).

Finally, we compared the epigenetic marks and expression profiles associated with different types of TEs, based on their methods of transposition. We noted distinct patterns of DNA methylation and histone modifications associated with class I and II TEs: class I TEs are enriched with CG methylation, co-localizing with or without CHH methylation, while class II TEs are predominantly marked by CHH DNA methylation, and CG and CHG methylation events co-localize with one another (Fig S9). A similar pattern was reported in soybean where a high abundance of CHH methylation over class II elements was correlated with the presence of small RNAs ^45^. This specific pattern might also be relevant to the nature of replication of each class of TEs; class I relies on a replicative mechanism for transposition while class II elements move by a cut and paste mechanism ^29^. Class I TE copy number can rapidly increase, and their higher methylation in both CG and CHG contexts might be a means to keep their expression under tight control. An interesting pattern of co-occurrence of epigenetic marks over TEs emerges from this analysis, which shows a systematic repression of TEs when associated with CHG methylation (Fig S9). Although the presence of CHG methylation is associated with active marks such as H3K9/14 Ac and H3K4me2, both classes of TEs show a decrease in their expression, suggesting the importance of maintenance of a heterochromatic environment. When checked, many of the TEs with CHG methylation were found to be inserted into or overlapping with genes (Table S8), reflecting the importance of maintaining these TEs in a silent state. A similar phenomenon has been observed for *Arabidopsis* TEs which are inserted into genes and whose repression is required to avoid the deleterious effects of TE insertion into host genes ^46^.

In summary the dissection of *P. tricornutum* genome reported here has led to the discovery of a significant number of novel genes with an important proportion of Rv genes with additional functional domains suggesting a role of Rv in restructuring genes and genomes during diatom evolution. Furthermore, our work brings new insights into the role of alternative splicing which we discovered to be abundant in *P. tricornutum* and is by no means restricted to respond to nitrate limitation and is likely to play a major role along with epigenetics in diatom response to environmental cues. Finally, our study contributes to better understand the complex chimeric nature of diatom genomes supporting the presence of large gene transfers from green lineages as well as red and bacterial genes. Overall, our data provides insights into the genetic and evolutionary factors that contribute to the ecological dominance and success of diatoms in contemporary oceans.

## Methods

### Data generation and mining

*Phaeodactylum tricornutum* genome re-annotation (named as Phatr3) was done on the Phatr2 genome assembly (ASM15095v2). The Phatr2 assembly was generated by the Joint Genome Institute (JGI), which resulted in 10,402 gene models from 33 assembled scaffolds (12 complete and 21 partial chromosomes) and 55 unassembled scaffolds ^7^. Gene models were predicted from RNA-Seq mapping and aligning the EST data-set using est2 genome. Additionally we used SNAP and Augustus and MAKER2 for final gene predictions. Apart from the previous assembly information, the species-specific data used in this re-annotation included the following.

#### RNA-Seq

Multiple RNAseq libraries (103 libraries in total) were generated under different growth conditions and are being used for many functional studies ^47^. The growth conditions used can be broadly divided into two major categories: 1) Nitrogen availability, which include 30 RNAseq libraries generated using Illumina platform (Bio-project accession no. PRJNA311568; Bio-sample accession numbers SAMN04488978-SAMN04489007), and 2) Iron availability, includes 49 libraries of RNA-Seq generated using SoLiD sequencing technology (SRA: SRP069841) ^47^. Apart from these 91 libraries, 12 more RNAseq libraries were generated including the wild type and alternative oxidase (Phatr2_bd1075) mutants ^48^; Bio-sample accessions: SAMN06350641-SAMN06350652. More information about the culture conditions can be referred from File S2.

#### Expressed sequence tags (ESTs)

Along with multiple RNAseq libraries existing 13,828 non-redundant *P. tricornutum* ESTs ^14,49^ done in different growth conditions were also utilized. Other EST data used includes 93,206 diatom ESTs from dbEST ^50^.

#### Epigenetic Marks

For better characterization of the genes both structurally and functionally, chromatin immunoprecipitation-sequencing (CHIPseq) data of multiple histone (H3) post-translational modification marks (H3K27me3; H3K9me2; H3K9me3; H3K4me2 and H3K9_14Ac) and DNA methylation data, from previous studies ^15,16,51^ were also included.

#### Construction of a multi-sequence reference dataset

A composite reference library, consisting of 75001602 non-redundant protein sequences was compiled from UniPROT (http://www.uniprot.org/help/uniref) (downloaded February 2016), alongside additional genomic and transcriptomic resources from JGI, MMETSP, and the 1kp project currently not located on UniPROT (http://genome.jgi.doe.gov; http://marinemicroeukaryotes.org/; https://sites.google.com/a/ualberta.ca/onekp/) (Table S4)^17^. To minimize artifacts arising from contamination between different MMETSP libraries, each MMETSP library was first pre-cleaned using a BLAST pipeline as described previously ^52^ which identifies a custom similarity threshold between each constituent library above which sequence pairs are inferred to be contaminants. In addition, following the methodology of a previous study ^10^, MMETSP libraries from a further twelve species were excluded due to the presence of larger scale systematic contamination (Table S4).

The reference sequence library was split into twenty-five prokaryotic sub-categories, including archaea and forty-nine eukaryotic sub-categories, which were finally binned into nine distinct groups ^10,19^ (Table S4; well described in File S2). These are diatoms, non-diatom stramenopiles, non-stramenopile SAR, CCTH (containing cryptomonads and haptophytes), green eukaryotes (including glaucophytes), red algae, amorpheans, prokaryotes and viruses. The taxonomic divisions were designed to reflect both current opinions regarding the global organization of the tree of life, as defined using up-to-date taxonomic information ^10,19^, and to provide enhanced resolution of the closest relatives of *P. tricornutum* (i.e. other diatoms, other stramenopiles, and other SAR clade members except for stramenopiles).

### Gene discovery and annotation

#### Structural re-annotation

The *P. tricornutum* version 2 genome (JGI), published in the year 2008 ^7^, was comprehensively structurally re-annotated using multiple RNA sequencing libraries, non-redundant ESTs and updated repertoires of stramenopile proteomes. 42 RNA-Seq libraries generated using an Illumina sequencing platform were mapped to the genome using Genomic Short-read Nucleotide Alignment Program, GSNAP ^53^, integrated within mapping pipeline of Ensembl Genomes ^54^. The remaining 49 SoLiD sequence libraries were aligned to the genome in color space ^55^. The alignment file for each mapped library can be accessed from ftp://ftp.ensemblgenomes.org/pub/misc_data/bam/protists/phatr3/. The mapping percentage was estimated using Samtools and varied between 70 – 96% (Table S9). Transcript assembly was further executed using Cufflinks ^56^ with default parameters. The resulting unfiltered transcripts along with EST libraries and protein sequences from stramenopiles UniProt Reference Clusters (UniRef90) ^57^ were then used for the genome re-annotation. The structural units derived were finally used to train SNAP ^58^ and Augustus ^59^ gene prediction programs using default parameters of the MAKER2 annotation pipeline ^60^.

#### Functional re-annotation

All predicted gene models were annotated for protein function using InterProScan ^61^ as part of the Ensembl protein features pipeline^62^. The results are available at http://protists.ensembl.org/Phaeodactylum_tricornutum/Info/Index/. We also used CLADE and DAMA to enhance the protein domain predictions using default parameters. From the 12177 genes, we succeed to predict the function and the domain architectures of 8235 by coupling CLADE and DAME predictions. For further 3942, we use only the best model output of CLADE. Although, CLADE predicted only one conserved region per gene, only 61 genes remain with unknown functions.

Next, the presence of organelle signaling signatures within the entire Phatr3 gene repertoire was further investigated using ASAFind and HECTAR, under the default conditions as specified in the original publications for each program ^63,64^. HECTAR was run remotely, using the Galaxy integrated server provided by the Roscoff Culture Collection (http://webtools.sb-roscoff.fr/).

#### Distribution of epigenetic marks

Data corresponding to histone H3 post-translation modification marks, H3K27me3; H3K9me2/3; H3K14_Ac; H3K4me2 and DNA methylation (CG, CHH, CHG), were taken from ^15^ and ^16,51^, respectively. RNA-Seq data for Pt18.6 (normal growth condition) was also downloaded from the same resource. Distribution of all marks along with the expression were then checked over the new genes and new transposable element (TE) models by adapting the methodology applied in ^15^ and ^16^. Estimation of expression (normalized DESeq counts) and differential gene expression analysis at different stages of the study was performed using Eoulsan ^65^, using parameters as indicated in Eoulsan parameter file used, (File S3). Statistical analysis to compare the expression of genes was performed using two-sample t-test with unequal variance.

#### RT-PCR

Total cellular RNA was extracted from approximately 30 ml late-log phase *P. tricornutum*, grown as described above, by phase extraction with Trizol (Thermo, France), followed by treatment using RNAse-free DNAse (Qiagen, France) and cleanup using an RNeasy column (Qiagen) as previously described ^10^. RNA was verified to be free of residual DNA contamination by PCR using previously generated universal 18S rDNA primers ^66^. cDNA was synthesized from 100 ng RNA-free DNA using a Maxima First cDNA synthesis kit (Thermo), and PCR was performed using the cDNA template and primers designed against the 5' and 3' ends of genes of interest using DreamTaq DNA polymerase (Thermo), per the manufacturers' instructions. Products were separated by electrophoresis on a 1%-agarose TAE gel containing 0.2 μg/ml ethidium bromide at 100V for 30 minutes, and visualized with a UV transilluminator. Representative products from each reaction were purified using PCR cleanup spin columns (Macherey-Nagel, France), and confirmed by Sanger sequencing (GATC, France) using both the forward and reverse PCR primers.

### Conservation analysis of Phatr3 gene repertoire

The evolution of the *P. tricornutum* genome was examined using gene homology searches. Orthologues of each gene were identified from each taxonomic sub-category, following the methodology used in the original *Phaeodactylum* genome annotation, by reciprocal BLAST best hit with an initial threshold e-value of 1 × 10^−10^. To minimize the effects of sequence contamination, and subgroup-specific gene transfer events, genes were only denoted as being shared with a particular group if reciprocal BLAST best in at least two separate taxonomic sub-categories within that group, following methodology established elsewhere ^10,67^.

### BLAST top hit analyses

A novel pipeline, based on BLAST top hit analysis, was designed to determine the probable gene transfer events that have occurred in the evolution of *P. tricornutum*, based on previous BLAST based reconstructions of gene sharing in other photosynthetic eukaryotes ^10,67^. For this analysis, the top BLAST hit from each taxonomic sub-category, for each gene in the *Phaeodactylum* genome, were collated to form a single reference library. To enable the identification of highly divergent or partial copies of each gene, a relaxed threshold e-value (10^−10^) that has previously been used for evolutionary analyses of diatom genomes, was employed ^7,11^. Following methodology established in a previous study ^10^ and the RbH analysis above, a gene was only deemed to have a particular evolutionary origin if top hits were obtained in two or more sub-categories within a particular lineage, prior to the best hit from another lineage. This analysis was performed for seven different reference libraries, one containing all possible reference sequences (Phatr3), and six deducting relatives of *P. tricornutum* (all pennate diatoms, all diatoms, all ochrophytes, all stramenopiles, all SAR clade members, and all SAR + CCTH clade members). Tabulated outputs for each analysis are provided in Table S3. The results obtained by this pipeline were compared to a subset of 324 Phat3 genes for which single-gene tree topologies generated using the expanded reference library have previously been published ^10^, and found to give broadly equivalent results (see Results; Fig. S10; Table S10).

Conceptual translations of the entire *Phaeodactylum* genome was searched against this modified library again using BLASTP, and the top ten hits for each gene were ranked. The group and sub-category for each BLAST top hit was profiled. BLAST top hits were only recorded if the top ten hits contained another sequence from a different sub-category within the same group, as defined using the taxonomic categories defined above, with a better expected value than the first hit from outside the same group as the top hit. For example, a BLAST output consisting of a first hit from a centric diatom, a second hit from a pennate diatom, and a third hit from a non-diatom group, would be considered to be genuine, whereas an output consisting of a first hit from a centric diatom, second hit from a non-diatom, and third hit from a pennate diatom would not. Genes for whom no BLAST hits were obtained were annotated as producing "no match". Genes for which top hits were identified, but were not taxonomically consistent with one another, as defined above, were annotated as being "ambiguous".

The BLAST top hit analysis was modified in two further ways, to allow more precise characterizations of different gene transfer events. First, the BLAST top hit analysis was repeated with five additional reference libraries, created using the same process as detailed above, but omitting six different groups of organisms, based on evolutionary proximity of the nuclear group to *P. tricornutum*: first, all pennate diatoms were removed, then all diatoms, then all ochrophytes, then all stramenopiles, then all SAR clade members, and finally all SAR and CCTH clade members ^10,19^. This was performed to allow inference of gene transfer events that have occurred in the ancient evolutionary history of *P. tricornutum*: for example, a gene that yielded a diatom top hit with the full library and pennate diatom-free libraries, a non-diatom stramenopile top hit in the diatom-free library, and a prokaryotic top hit in the stramenopile-free library would be inferred to have been undergone a lateral transfer between prokaryotes, and an early ancestor of stramenopiles, and to have been vertically inherited by *Phaeodactylum tricornutum* from the stramenopile ancestor onwards. Secondly, all of the BLAST top hit analyses were repeated using modified libraries from which all groups with a suspected history of secondary endosymbiosis (i.e. cryptomonads, haptophytes, myzozoans, chlorarachniophytes, and euglenids) ^10^.

### Alternative splicing

To explore the set of genes undergoing alternative splicing, exon skipping or intron retention, 17 RNA-Seq samples prepared under different conditions of nutrient availability (Biosample accessions: SAMN06350643, SAMN06350647, SAMN04488984, SAMN04488988, SAMN04488978, SAMN04488992, SAMN04488980, SAMN04488981, SAMN04488985, SAMN04488979, SAMN04488989, SAMN04488983, SAMN04488987, SAMN04488991, SAMN04488982, SAMN04488986, SAMN04488990) were compared. Only genes that were annotated as having two or more exons, and containing introns with a minimum length of 50 bp, were considered for the analysis. RNA-Seq reads were mapped on the reference genome using Bowtie ^68^ with parameters: -n 2 -k 2 --best. To filter the significant candidate features, we considered horizontal (along the gene) and vertical coverage (depth of reads) to be more than 80% and 4x, respectively. Theoretical support to the candidate features showing exon-skipping or intron-retention was provided by measuring the rate of consensus observation within multiple samples studied. For exons anticipated to show exon-skipping, the observation had a consensus from more than 20% and less than 80% samples. On the other hand, introns having a consensus observation of their retention from more than 20% samples were considered as true events. Functional association studies were further performed on genes showing evidence for exon-skipping or intron-retention. Genes were clustered based on conditions used to prepare the RNA-Seq samples and gene ontology (GO) terms were assigned to the genes (wherever possible) using UniProt-GOA (http://www.ebi.ac.uk/GOA). Significance of these terms was interpreted by calculating the observed to expected ratio of their percent occurring enrichment. The occurrence of an individual biological process within a specific functional set (genes exhibiting intron-retention/exon-skipping, etc.) was compared to that of its occurrence in the complete annotated Phatr3 biological process catalog. The degree of significance of enrichment of each biological process was quantified using a chi-squared test, with a threshold significance P value of 0.05.

To gain insights into the role of alternate splicing in regulating the molecular physiology of the cells, the expression patterns of AS candidates predicted to undergo IR and/or ES under nitrogen replete conditions (Biosample accessions: SAMN04488981, SAMN04488985, SAMN04488979, SAMN04488989) were compared at different time-points (T15min, T45min, T90min and T18hrs (referred as Tend) to the wild-type (Biosample accession: SAMN06350643). RNA-Seq expression values were calculated using Eoulsan ^65^, and additionally normalized to eliminate biases caused by the restructuring of alternatively spliced transcripts.

### Annotation of repetitive elements

The REPET v2.2 package ^69^ was used to detect the repetitive fraction of the *P. tricornutum* genome. The TEdenovo pipeline (https://urgi.versailles.inra.fr/Tools/REPET) was launched including the Repeat Scout approach ^70^ to build a library of consensus sequences representatives of the repeated elements in the genome assembly. The classification comes from decision rules applied to the evidence collected from the consensus sequences. These include: search for structural features, search for tandem repeats, comparison to PFAM, comparison to Repbase ^71^ and to a local library of known TEs. The library of manually curated TEs that has been established in previous work ^44^ as appended to the TE denovo library and redundancy was removed from the combined library. The TE annotation pipeline was then run with default settings using the sequences from the filtered combined library as probes.

## Acknowledgements and Funding

We thank Stephane Le Crom and Laurent Jourdren for helping setting up EOULSAN. CB acknowledges funding from the ERC Advanced Award ‘‘Diatomite’’, the Louis D Foundation of the Institut de France, the Gordon and Betty Moore Foundation, and the French Government ‘‘Investissements d’Avenir’’ programmes MEMO LIFE (ANR-10-LABX-54), PSL* Research University (ANR-1253 11-IDEX-0001-02), and OCEANOMICS (ANR-11-BTBR-0008). CB also thanks the Radcliffe Institute of Advanced Study at Harvard University for a scholars fellowship during the 2016-2017 academic year. AR was supported by a MEMO LIFE PhD scholarship and RGD was supported by an EMBO Long-Term Fellowship (ALTF 1124-2014).

## Supplementary Figures

**Figure S1.**
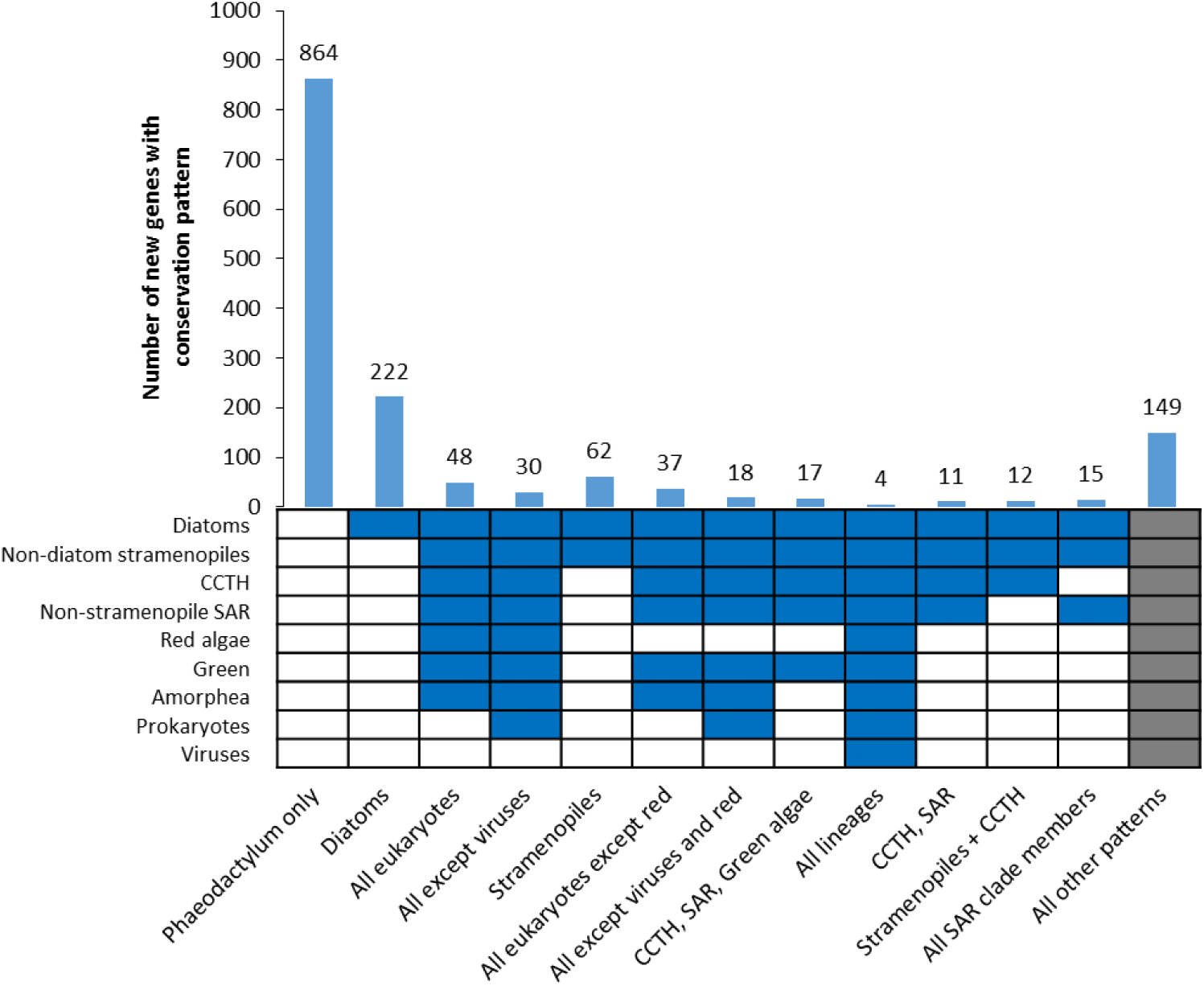
Numbers of novel genes in Phatr3 identified as being shared with different groups of organisms. The heatmap and graph are shown as per Fig 1.

**Fig. S2.**
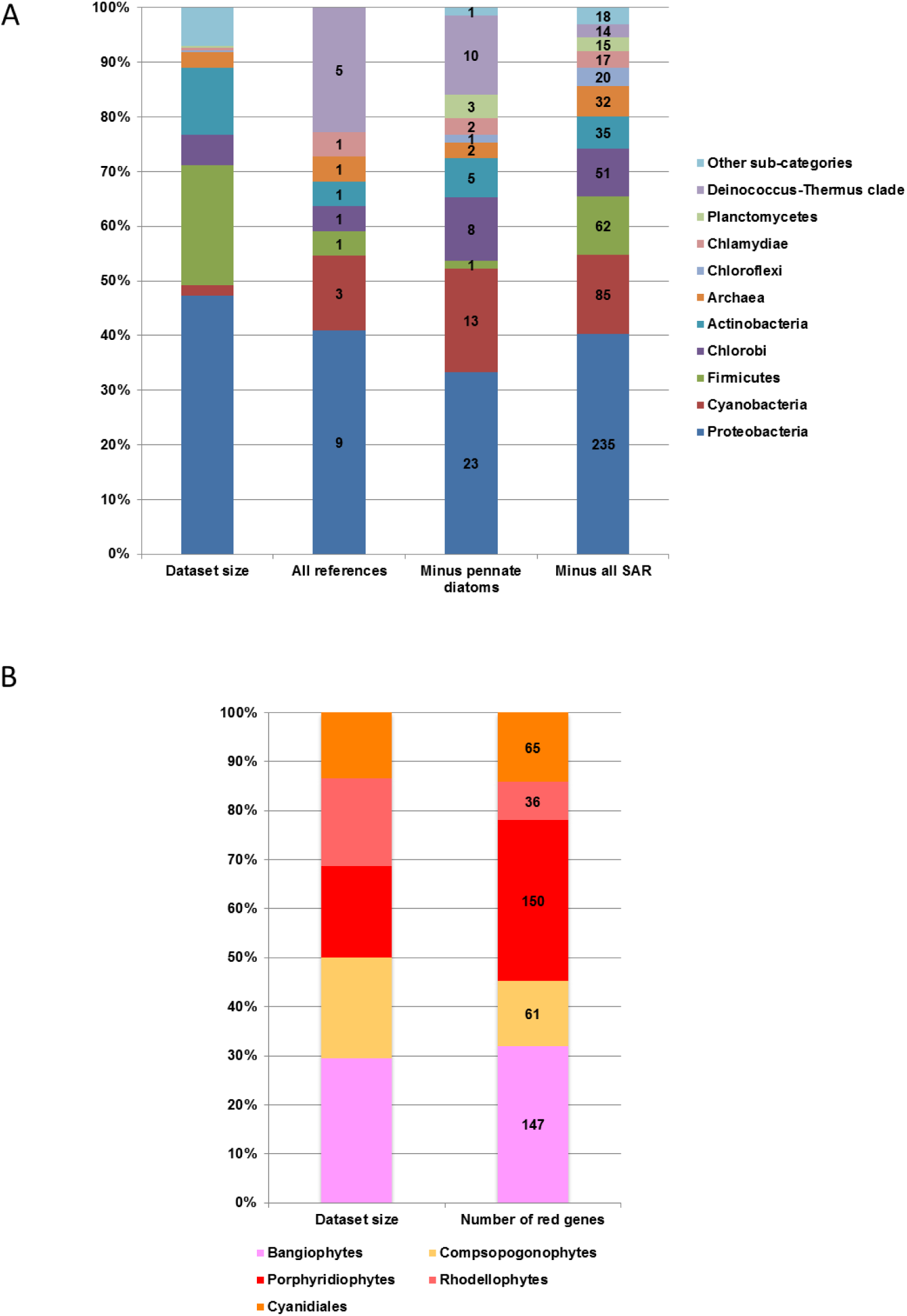
Specific taxonomic affiliations of Prokaryotic, Red and Green genes. Each chart shows the specific sub-category from which different prokaryotic (A), and red (B) genes arose. All genes that were assigned (i.e., two or more top hits from two or more sub-categories from a particular lineage, prior to the first top hit from outside that lineage) using the most reduced reference dataset (i.e., all reference sequences, excluding SAR clade members, and other algal lineages with secondary or tertiary plastids) is shown. For prokaryotic genes, two other distributions (obtained for the entire dataset minus non-ochrophyte algae with secondary or tertiary plastids, and the entire dataset minus pennate diatoms, and all non-ochrophyte algae with secondary or tertiary plastids) are shown. Each chart additionally shows the relative size of each sub-category within the reference sequence library, demonstrating that certain sub-categories contribute to substantially more of the top hits (e.g., the *Deinococcus-Thermus* clade, in the distribution of prokaryotic genes for the full and pennate diatom-free datasets that were modified to remove all non-ochrophyte lineages with secondary or tertiary plastids) or fewer of the top hits (e.g., the streptophytes, in the distribution of green genes for the dataset from which all SAR clade sequences, and other non-ochrophyte lineages with secondary or tertiary plastids were removed) than might be expected given the corresponding dataset size.

**Fig. S3.**
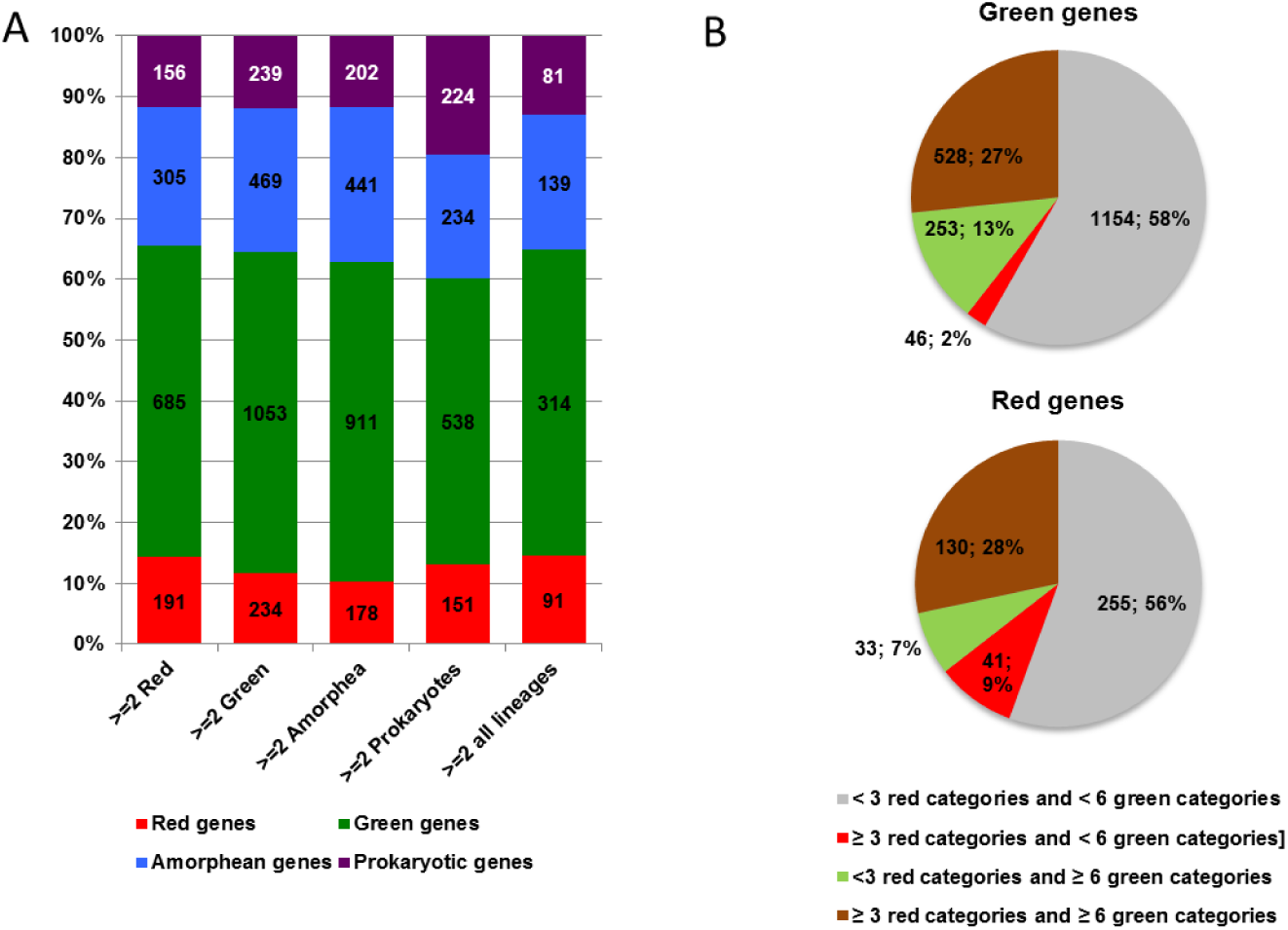
Green genes are not purely a result of taxonomic undersampling or lineage-specific gene loss. (A) shows the taxonomic affiliations of genes identified by BLAST top hit analysis, with the dataset from which all SAR clade sequences, and other non-ochrophyte lineages with secondary or tertiary plastids were removed, for which orthologues could be identified in at least two red, green, amorphean or prokaryotic sub-categories, and for which orthologues could be identified in at least two each of the red, green, amorphean and prokaryotic sub-categories. In each case, substantially more genes of green affinity were identified than of other taxonomic affiliation. (B) Compares the number of genes of red or green taxonomic affiliation for which RbH orthologues could be identified in a majority of red (3/5) or green (6/11) sub-categories. A similar proportion of genes of inferred red origin (130/459, 28%) and genes of inferred green origin (528/ 1981, 27%) were found to have orthologues in a majority of both red and green sub-categories, indicating that the identification of green genes within the dataset was not unfairly biased by taxonomic undersampling of red lineages.

**Figure S4.**
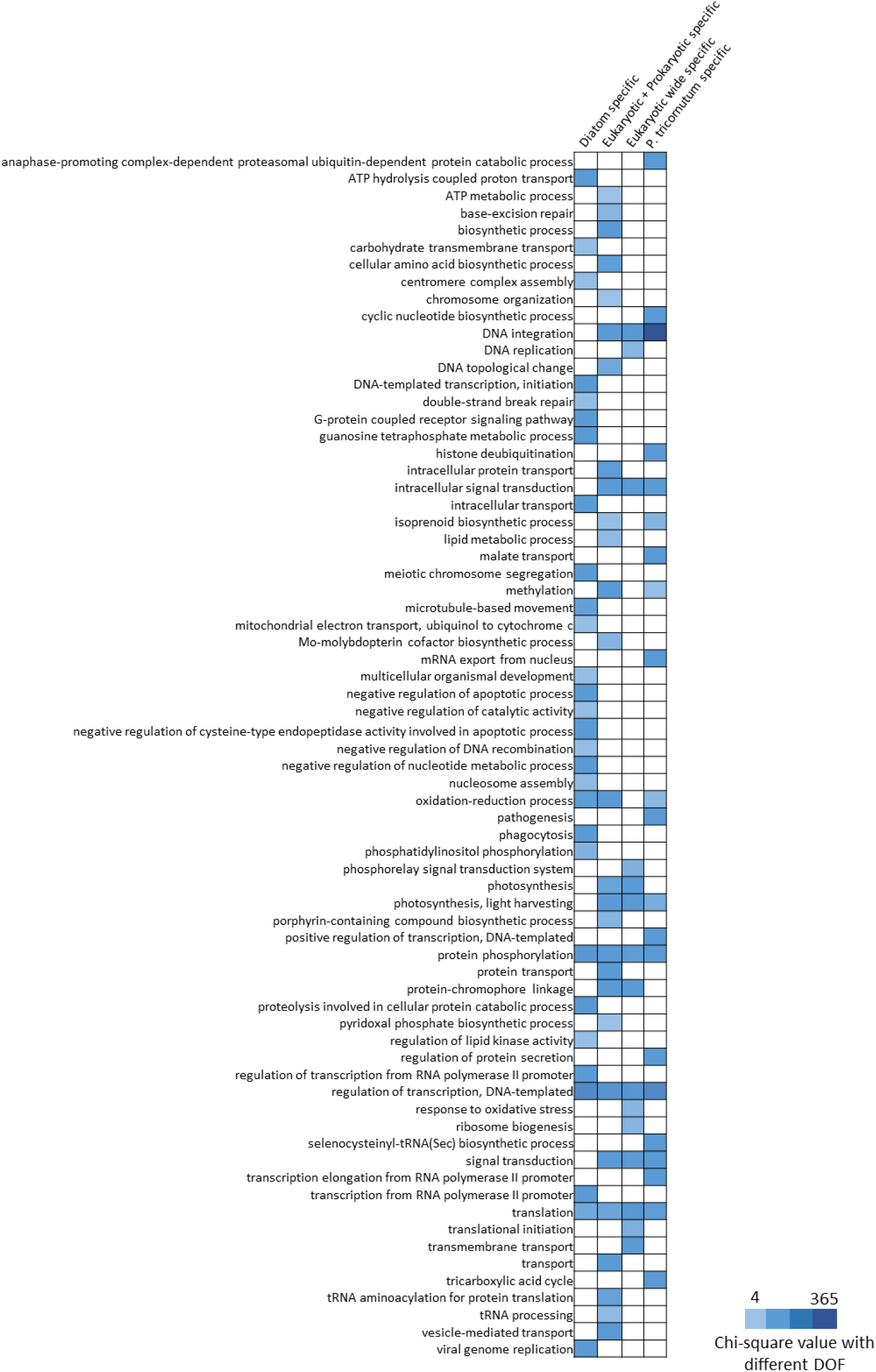
Enrichment of biological processes within genes identified to be specific to different groups of organisms. The heat map, indicating chi-square values which are significant (P-value < 0.05) with different degrees of freedom (DOF), depicts various biological processes (left Y-axis) that are enriched in the pool of genes found specific to different groups of organisms (top X-axis). Chi-square values are used to rank the most significant biological processes in descending order. High chi-square value here indicates higher significance (very low P-value) compared to low chi-square values indicating higher P-value but < 0.05.

**Fig. S5.**
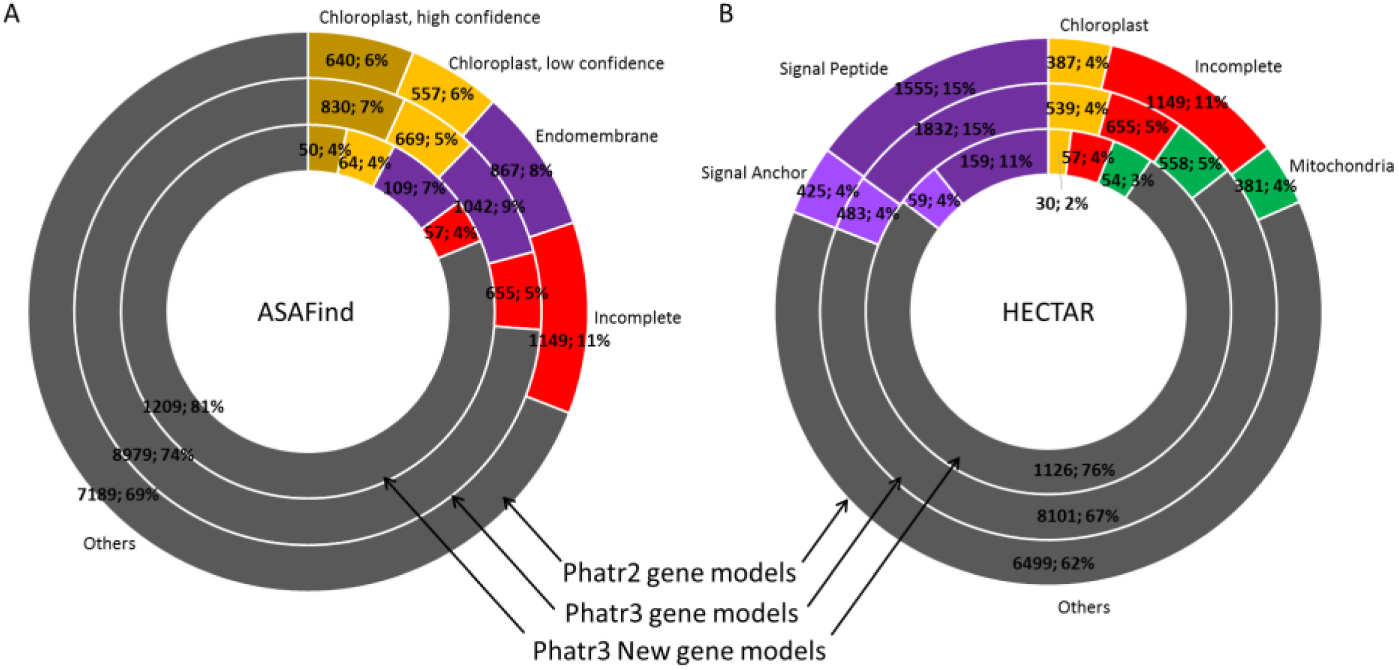
Predicted subcellular localization of *P. tricornutum* proteins. This figure shows the targeting predictions for proteins encoded within the *P. tricornutum* genome as assessed using the diatom targeting predictor programmes (A) ASAFind (Gruber et al., 2015) and (B) HECTAR (Gschoessl et al., 2008). The figure is in accordance with Figure 3 panel A and B.

**Figure S6.**
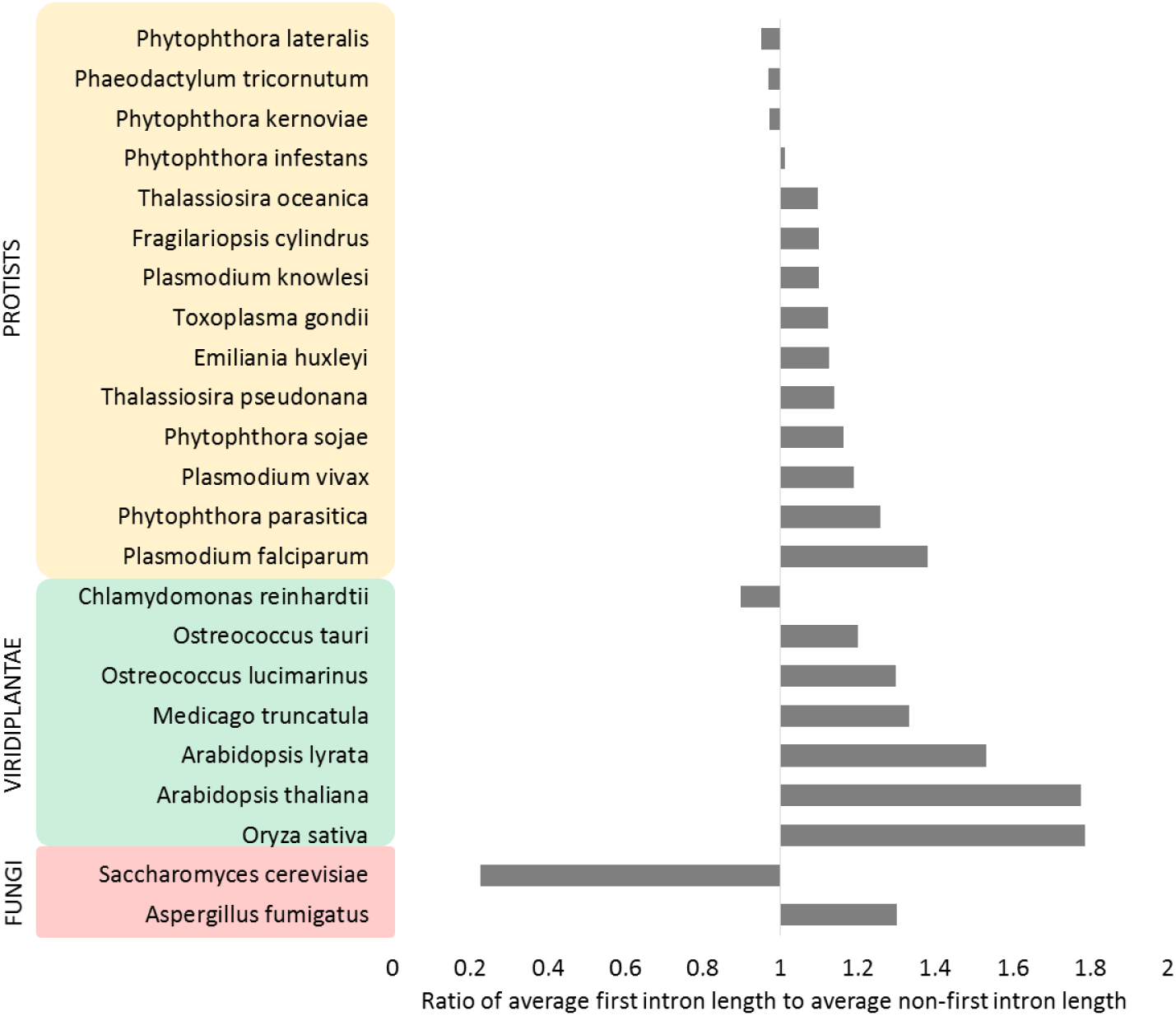
Comparative analysis of first intron length to non-first intron length. The bar-plot represents the comparative meta-gene analysis of ratio of average first intron length to average non-first intron length between multiple Protists, Plants and Fungal species. Annotation data was taken from Ensembl.

**Figure S7.**
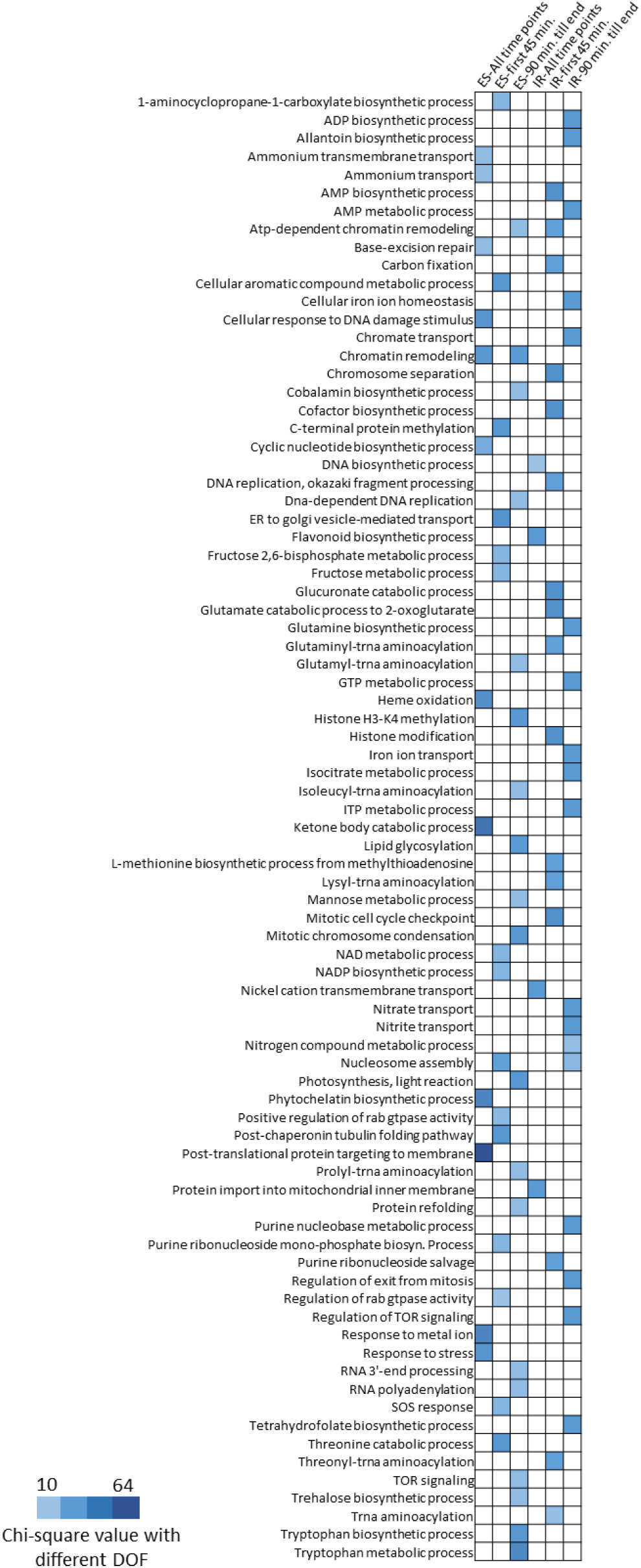
Enrichment of biological processes within genes exhibiting alternative splicing at various time-points under Nfree culture conditions. The heat map, indicating chi-square values which are significant (P-value < 0.05) with different degrees of freedom, depicts various biological processes (left Y-axis) that are enriched in the pool of genes exhibiting alternative splicing in the context of intron-retention and exon-skipping (top X-axis). The figure is in relation with categories represented in Figure 4 panel C. Chi-square values are used to rank the most significant biological processes in descending order. High chi-square value here indicates higher significance (very low P-value) compared to low chi-square values indicating higher P-value but < 0.05.

**Figure S8.**
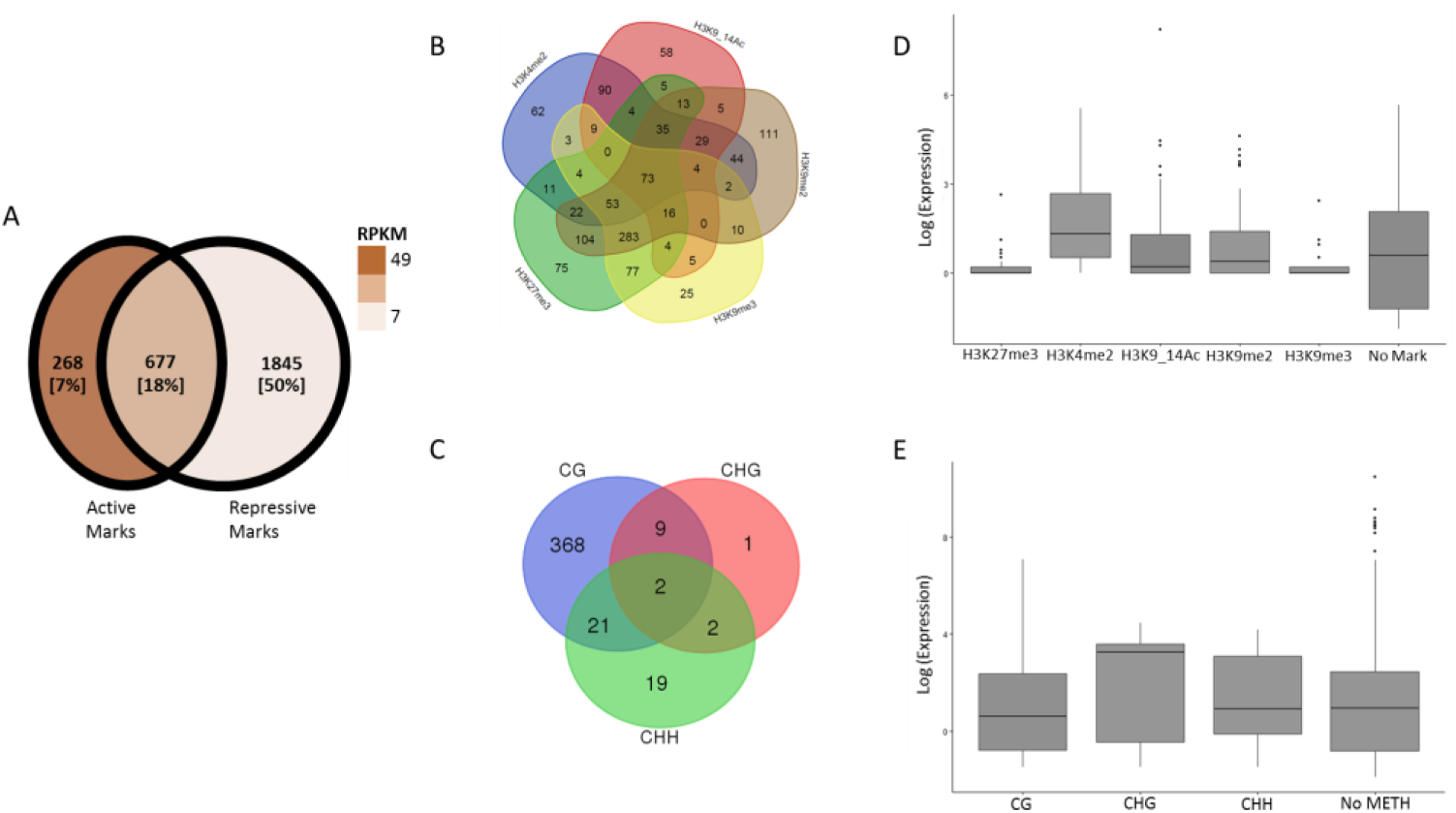
Distribution of epigenetic modifications over transposable elements. The Venn diagram (A) represents different chromatin states maintained based on the association of TEs with repressive, active or both repressive and active chromatin modifiers. Numbers and percentages in the Venn diagram reflects the absolute number of TEs and the relative percentage out of the total Phatr3 TEs. The Venn diagram in (B) presents the number of new TEs found to be localized by one or more histone H3 PTMs, and (C) presents the new TEs methylated in different context (CG, CHH, and CHG) of DNA methylation. Boxplots (D) and (E) represents average (median) expression of genes marked exclusively either of the H3 PTMs and are DNA methylated in either of the context, respectively, in normal condition.

**Figure S9.**
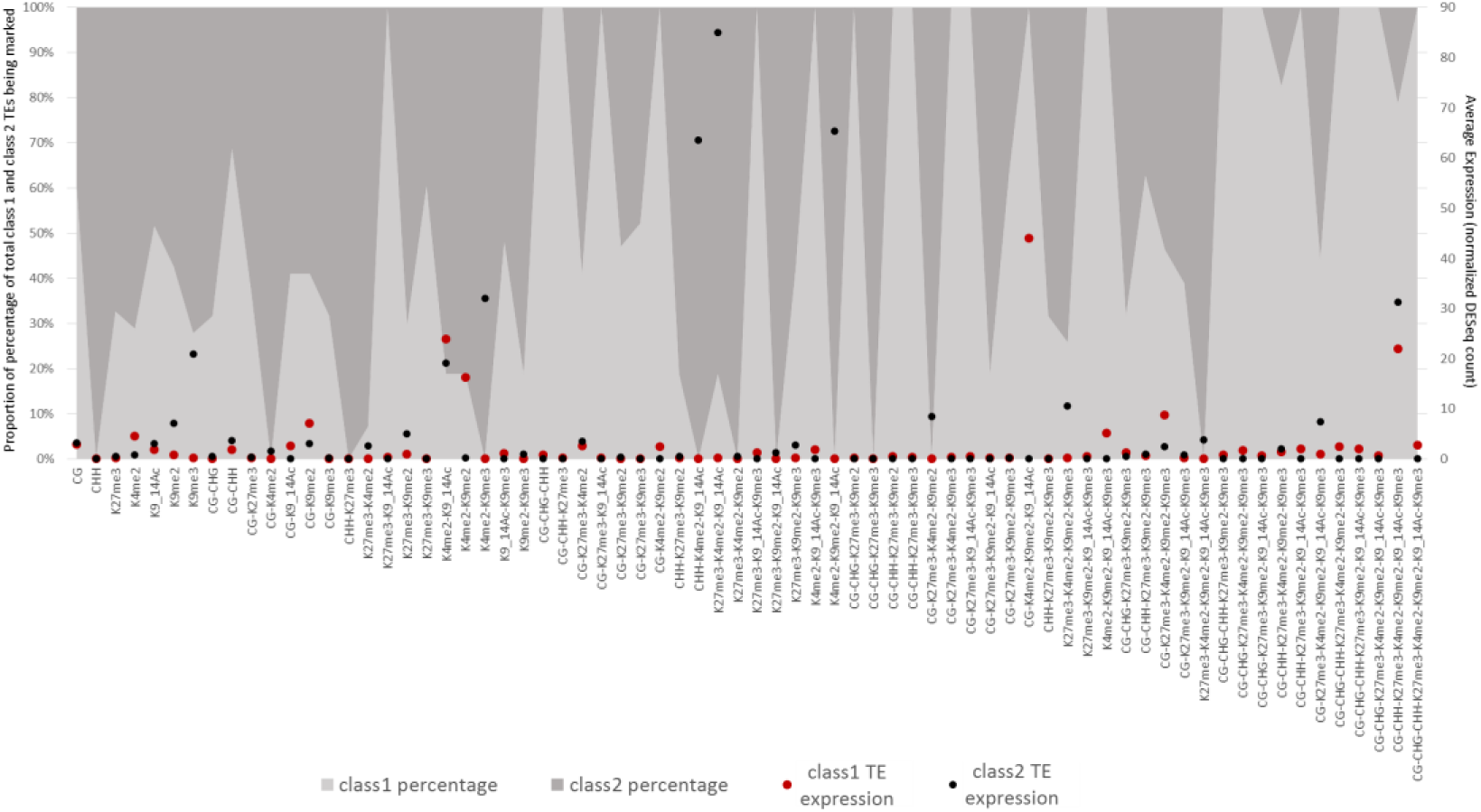
Epigenetic marking over transposable elements. The area plot represents the proportion of Class I vs Class II transposable elements being marked by different epigenetic marks including Histone H3 post-translational modifications and DNA methylation (CG, CHH and CHG). Black and red dots indicate the average RNA expression of all the TEs (wherever available) marked in different contexts.

**Fig S10.**
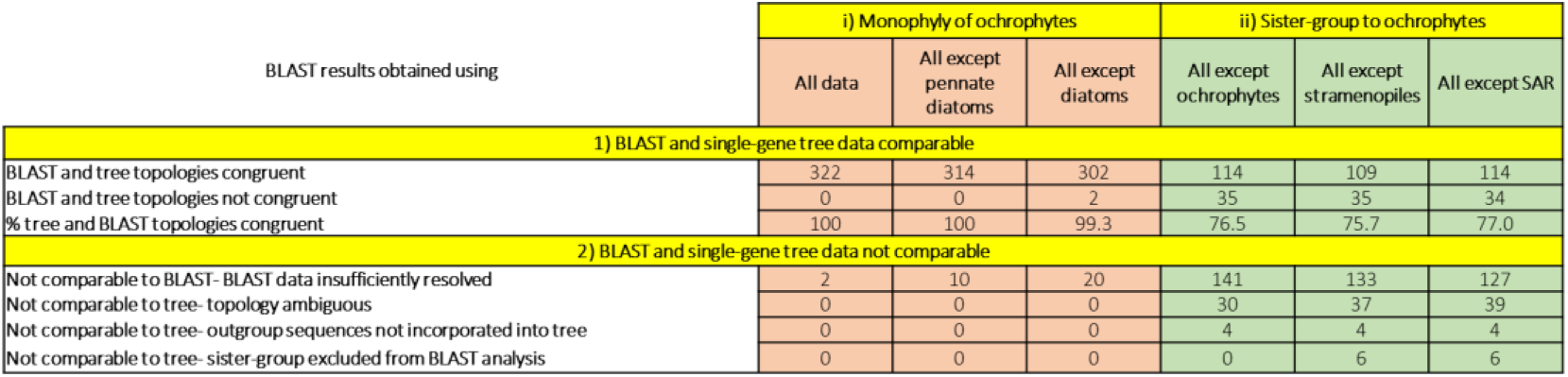
Verification of the reconstruction of evolutionary origins by BLAST top hit analysis. **(A)** Compares the results of BLAST top hit analysis and single-gene phylogeny for 324 genes in Phatr3 incorporated into an independent phylogenetic study of plastid-targeted proteins with broad ochrophyte distribution ^10^. Each of the proteins incorporated are found to produce a monophyletic or paraphyletic ochrophyte clade, i.e., should produce BLAST top hits to diatom or other ochrophyte sub-categories in the raw BLAST top hit analysis, and in BLAST top hit analyses from which pennate diatoms and all diatoms have been removed (but other ochrophytes have been retained). In addition, each protein should have a similar BLAST top hit in analyses from which all ochrophyte, stramenopile or SAR clade sequences have been removed to the sister-group to the ochrophyte clade (either red algae, green algae, aplastidic stramenopiles, other eukaryotic lineages, or prokaryotes) inferred from the single-gene tree. The overwhelming majority of the BLAST top hit analyses support monophyly of the ochrophytes, and at least three quarters retrieve the same ochrophyte sister-group as determined through single-gene tree analysis.

